# A diverse and distinct microbiome inside living trees

**DOI:** 10.1101/2024.05.30.596553

**Authors:** Wyatt Arnold, Jonathan Gewirtzman, Peter A. Raymond, Marlyse Duguid, Craig Brodersen, Cade Brown, Naomi Norbraten, Qespi T’ika Vizcarra Wood, Mark A. Bradford, Jordan Peccia

## Abstract

Despite significant advances in microbiome research across various environments^1^, the microbiome of Earth’s largest biomass reservoir– the wood of living trees^2^– remains largely unexplored. This oversight neglects a critical aspect of global biodiversity and potentially key players in tree health and forest ecosystem functions. Here we illuminate the microbiome inhabiting and adapted to wood, and further specialized to individual host species. We demonstrate that a single tree can host approximately a trillion microbes in its aboveground internal tissues, with microbial communities partitioned between heartwood and sapwood, each maintaining a distinct microbiome with minimal similarity to other plant tissues or nearby ecosystem components. Notably, the heartwood microbiome emerges as a unique ecological niche, distinguished in part by endemic archaea and anaerobic bacteria that drive consequential biogeochemical processes. Our research supports the emerging idea of a plant as a “holobiont”^3,4^—a single ecological unit comprising host and associated microorganisms—and parallels human microbiome research in its implications for host health, disease, and functionality^5^. By mapping the structure, composition, and potential sources and functions of the tree internal microbiome, our findings pave the way for novel insights into tree physiology and forest ecology, and establish a new frontier in environmental microbiology.

## Introduction

Trees are foundational components of the terrestrial biosphere and play a critical role in shaping ecosystem structure ^6^, stability ^7^ and biodiversity ^8^. Trees serve as hosts for other plants ^9^ and animals ^10^, regulating ecosystem services such as water filtration and nutrient retention ^11^, providing innumerable social and cultural values ^12^, and performing carbon fixation and sequestration ^13^, storing over 300 gigatons of carbon globally ^14^, primarily in wood ^15^. Wood underpins a vast array of economic activities including food, fuel, and fiber production and has an estimated global value of $9 trillion ^16^. Elucidating the biophysiochemical factors driving tree and forest health, function, and distribution are crucial to understanding Earth’s ecosystems, forecasting forest response to future change, and promoting ecological and economic security.

The field of microbiome research has revealed the profound and varied effects that consortia of bacteria, fungi, and archaea can have on the well-being of their host ^5^. In human biology and medicine, extensive investigations continue to unravel the implications of microbial diversity for both wellness and disease ^17^. Concurrently, environmental microbiome research has explored the structure and function of free-living and host-associated microbial communities across diverse environments (particularly animal guts, soils, and plant surfaces) ^1,18,19^. Comparatively little inquiry has gone into characterizing the microbial constituents inhabiting the largest reservoir of biomass on Earth ^2,15^: the wood of living trees.

The study of tree-associated microbiomes has focused principally on exposed surfaces ^20^, such as the roots ^21^, leaves ^22^, and bark of trees ^23^, rather than their wood ^24^, despite wood composing the vast majority of tree biomass. Whereas the existence of internal (“endophytic”) microbiota in plants has long been recognized ^25–30^, and the endophytic microbial community has been explored in model herbaceous and agricultural plants ^31^, microbiome studies of the dominant component of tree biomass, the boles of living trees, is limited. This is surprising given the long-recognized and profound impacts that microbial actors living in wood can induce, including enhanced growth ^32^ or disease and mortality. Outbreaks of fungal and bacterial pathogens, such as *Ophiostoma ulmi* ^33^ or *Xylella fastidiosa* ^34^, both of which target the wood of living trees, have rapidly devastated ecologically, culturally and economically important tree species in recent history.

In this study, we surveyed the wood-associated microbiome in more than 150 living trees across 16 species from the northeastern United States, representing 11 genera which have a global distribution across 151 countries (**SI Figure 1**). Using newly developed, novel sample processing methods ^35^, and quantitative and qualitative characterizations of microbiomes, we evaluated how wood-borne communities—both prokaryotic and fungal—vary within and among tree species, and how these consortia relate across a tree’s tissues and with neighboring environments. Through this microbial census, we illuminate the unexplored microbial biodiversity in the wood of living trees and provide insights into future forest health and ecosystem dynamics, which may lead to innovative strategies to conserve and enhance the vital functions of trees in ways that support global ecosystems and the communities that rely on them.

### The Wood of Living Trees Harbors a Quantitatively Substantial Microbiome

We estimate that there is approximately 1 bacterial cell for every 20 plant cells within the wood of the average living tree (based on observed prokaryotic abundances, which averaged 10^6^.^35^ 16S rRNA copies per dry gram wood). For the human microbiome, the ratio of bacterial cells to human cells is closer to parity (∼1:1) For a tree weighing approximately 5000 kg (live mass), the number of prokaryotes in the wood alone (notably, excluding the rhizosphere and external surfaces) is approximately 1 trillion cells, or one to two orders of magnitude less the number resident in the average human ^36^. This fractional number is in line with expectations, as wood possesses no sites akin to the mammalian intestinal tract, wherein labile substrates abound. Wood itself consists primarily of cellulose and lignin, which are less readily degraded by bacteria ^37^, along with nonstructural carbohydrates (NSCs), a potential energy source for woodborne prokaryotic life (and which constitute 5-10% of a trees’ stem by dry mass) ^38^. These NSCs are a collection of mobile sugars and starches used in metabolic functions, and are a recognized microbial substrate in the rhizosphere ^39^. However, living trees also contain many known antimicrobial compounds, such as phenolics produced by the living tissue embedded in the xylem ^40,41^, likely restricting runaway microbial growth. As such, the wood of living trees may reasonably be compared to an environment like the human stomach, wherein a balance between substrate availability and the antimicrobial action of stomach acid limit cell counts to 10^3^-10^4^ cells/mL ^36^.

For 11 of the 16 species studied, there was no significant difference in prokaryotic abundance between heartwood and sapwood (Wilcoxon, p>0.05) tissues. As we examined a range of species with variable patterns of heartwood formation, we operationally defined “heartwood” as the innermost 5 cm of wood tissue and “sapwood” as the outermost 5 cm of wood tissue ^35^. Despite an overall lack of difference between tissues, two species, *Acer saccharum* (red maple) and *Betula lenta* (black birch) displayed significantly higher prokaryotic abundances in their heartwood, whereas three species, *Fraxinus americana* (white ash), *Pinus strobus* (Eastern white pine), and *Quercus rubra* (red oak) displayed significantly higher prokaryotic abundances in their sapwood (p<0.05). The highest average (sapwood and heartwood) prokaryotic abundances, both near 10^7^.^3^ cells dry g^−1^ (**Figure 1**), were observed in the maples (*Acer spp.*), a group well known for abundant non-structural carbohydrates ^42^, whereas the lowest average abundances, at 10^4^.^8^ and 10^5^.^7^ dry g^−1^, were observed for *Prunus serotina* (black cherry) and *Tsuga canadensis* (Eastern hemlock), both of which possess abundant known antimicrobial compounds ^43,44^. Whether differing abundances between species result from active biological regulation or incidental habitat differences, these observations begin to portray distinct physical and biochemical microenvironments within trees as potential controlling factors of microbial growth and diversification.

**Figure 1:**
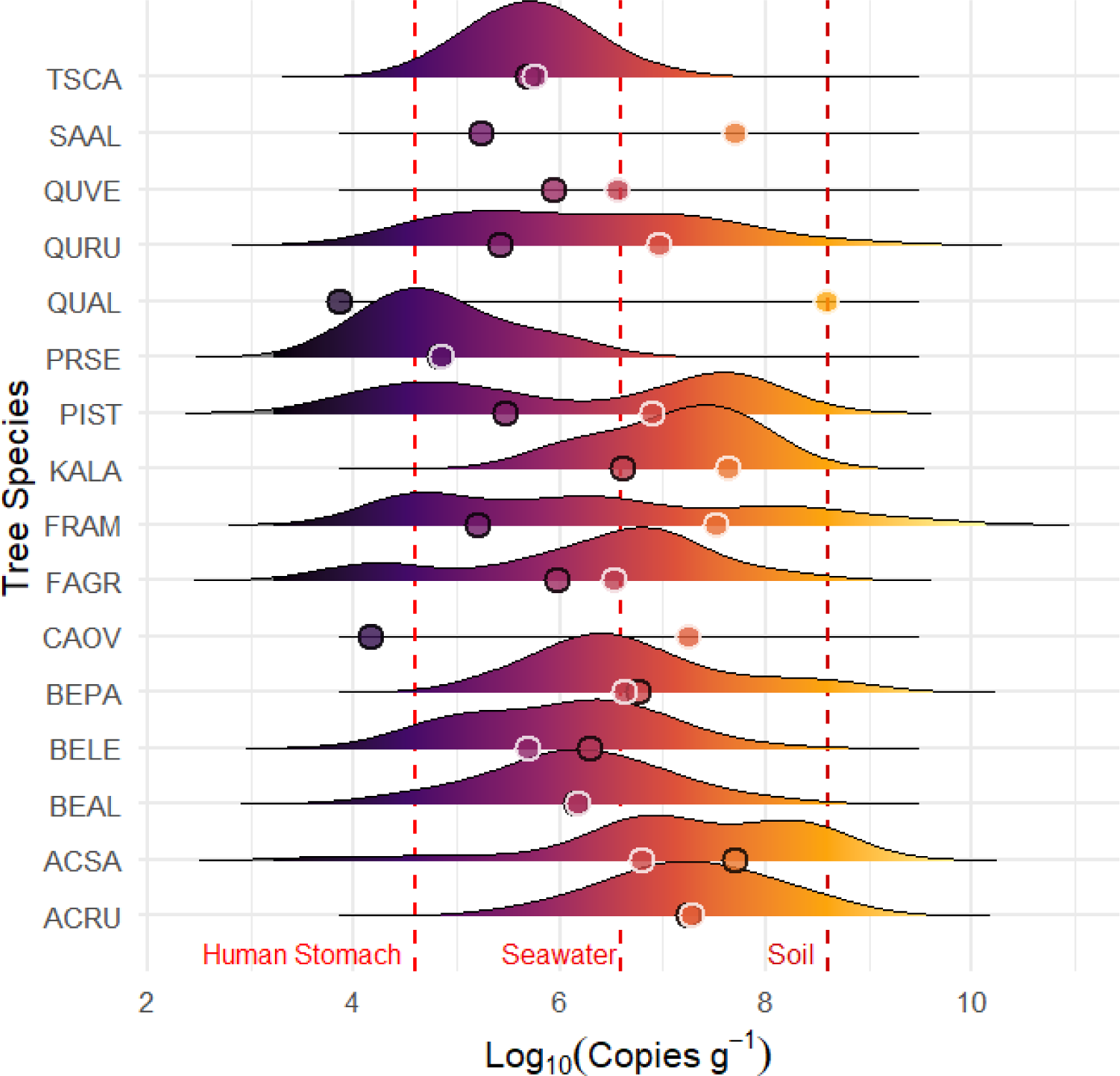
Prokaryotic abundances within the wood of living trees, as determined by 16S qPCR (per dry gram wood). Abundance distributions are split by species (for species codes, see **SI Table 1**), with the mean heartwood abundance identified in black, and the mean sapwood abundance in white. Estimates of the average 16S copies (assuming 4 copies per cell ^45^) in three different biological media (per gram of human stomach fluid ^36^, seawater ^46^, and soil ^47^) are identified in vertical red dashed lines. The number of individuals sampled per species ranged from n=1 to n=20 (see Materials and Methods); a distribution is not shown for species with n<3.

### A Consequential Mix of Environmental and Endemic Taxa

The wood of living trees included generalist groups common to other natural environments, but also a suite of taxa potentially endemic to the niche of tree boles. For instance, at the class and phylum levels, average abundances of Gammaproteobacteria, Alphaproteobacteria, and Actinobacteria constituted 9.1-29.3% of wood taxa, with these same taxa comprising sizable (7.2-19.5%) portions of the concurrently sampled soil microbiome. However, the wood also included high abundances (up to 13.5%) of distinct classes, such as Clostridia, Bacilli, Negativicutes, and Methanobacteria, which were found at <1% abundance in neighboring soil samples (**Figure 2**). Some of these unique groups, like those within the phyla Actinobacteria and Firmicutes, contain recognized, dominant plant endophytes, though observations of endophytic archaeal groups, like the aforementioned Methanobacteria, are less common in the literature ^48^. These observations suggest that living wood communities may share overlaps with better studied model plants and agricultural crops, but differ in new and substantial ways.

**Figure 2:**
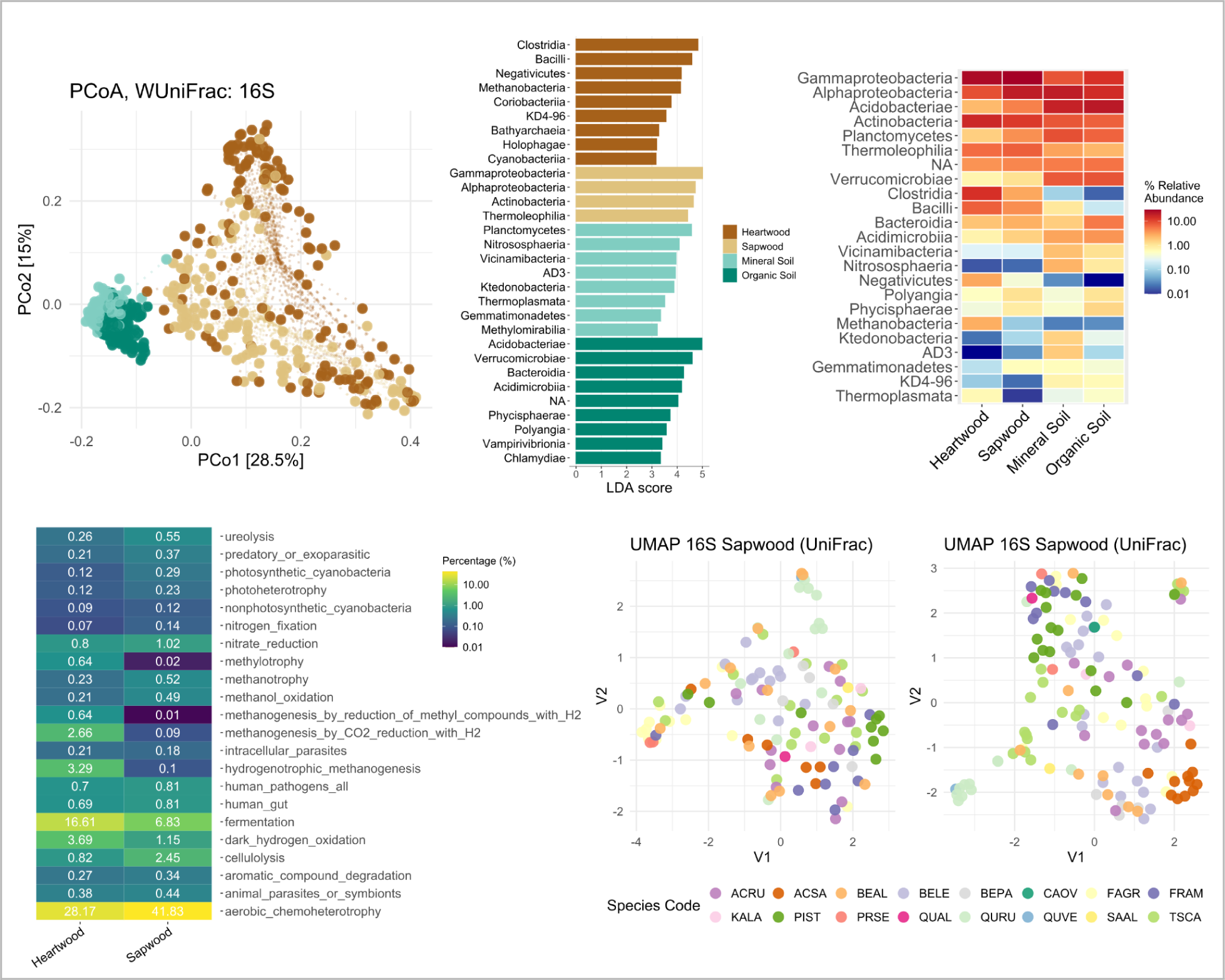
Bacterial and archaeal assemblages of the woodborne microbiome. **a.)** Principal coordinate analysis of 16S taxa using weighted UniFrac distance displays differentiation between wood samples (brown) and soil samples (teal). Dashed lines converge on sample type centroid. **b.)** Differential abundance analysis between wood (heartwood, sapwood) and soil (organic, mineral fractions) using LEfSe (p<0.01, LDA threshold set at ≥ 3.5). **c.)** Heatmap of relative abundances of prokaryotic taxa at the class level across the four sample types (top 23 taxa by abundance; “NA” refers to taxa unresolved at the class level). **d.)** Heatmap of prokaryotic metabolisms within different wood tissues, as inferred by FAPROTAX (percentages abundance weighted, metabolisms less abundant than 0.05% excluded). UMAP ordinations of **e.)** 16S sapwood and **f.)** 16S heartwood communities based on unweighted UniFrac distance, colored by tree species.

For fungi, a similar trend was observed, as classes such as Agaricomycetes, Leotiomycetes, and Sordariomycetes constituted large portions (3.8-48.7%) of both the wood and soil microbiomes (**Figure 3**). Interestingly, fewer unique classes were apparent for fungi, save for the notable exception of lichen (Lecanoromycetes), though this may have been a function of a high number of unresolved taxa. The abundance of saprotrophic groups, like Agaricomycetes, within the wood of living trees is in alignment with our existing understanding of endophytic fungi as a source of deadwood colonization and decay ^49^. Although their myriad roles in the wood of living trees remains a topic of active research ^50–55^ the presence of taxa like Agaricomycetes corroborates that trees are primed for colonization by their own endophytes upon death ^56^.

**Figure 3:**
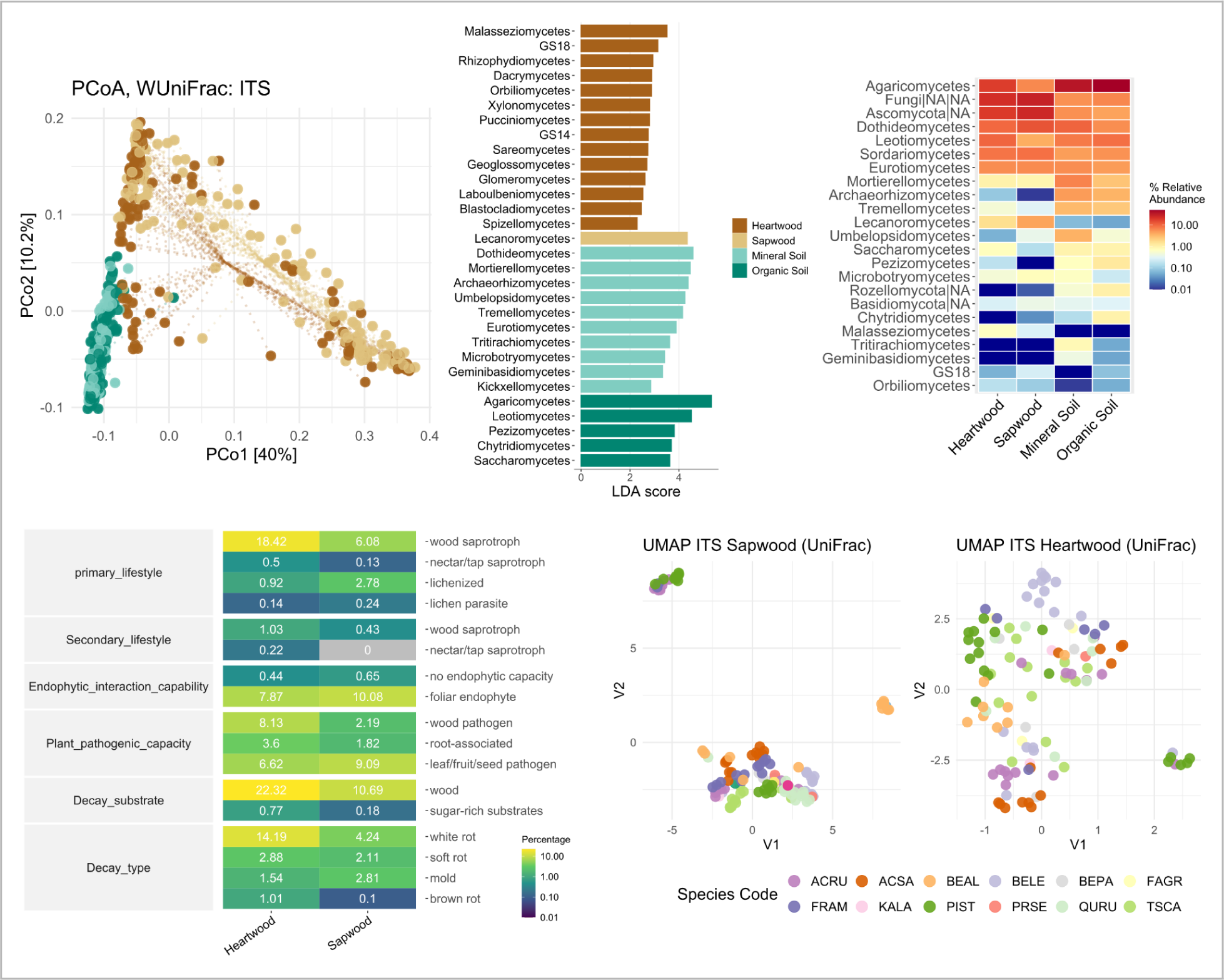
Fungal assemblages of the woodborne microbiome. **a.)** Principal coordinate analysis of ITS taxa using weighted UniFrac distance displays differentiation between wood samples (brown) and soil samples (teal). Dashed lines converge on sample type centroid. **b.)** Differential abundance analysis between wood (heartwood, sapwood) and soil (organic, mineral fractions) using LEfSe (p<0.01, LDA threshold set at ≥ 3.5). **c.)** Heatmap of relative abundances of fungal taxa at the class level across the four sample types (top 23 taxa by abundance; “NA” refers to taxa unresolved at the kingdom|phylum|class level). **d.)** Heatmap of prokaryotic metabolisms within different wood tissues, as inferred by FungalTraits (percentages abundance weighted, metabolisms less abundant than 0.05% excluded). UMAP ordinations of **e.)** ITS sapwood and **f.)** ITS heartwood communities based on unweighted UniFrac distance, colored by tree species.

Such insights into the likely candidates to initiate wood decay upon tree death, may also hold for prokaryotes ^57^. For instance, the abundance of Firmicutes in decaying wood is known to decrease as wood age increases ^58^. As Firmicutes, including Clostridia and Bacilli, are some of the most abundant members of the herein observed wood microbiome, we may reasonably expect their successional replacement following tree mortality and over the course of decay, as the conditions which once favored their presence (e.g., low oxygen content) shift following tree death ^59^.

As taxonomic resolution increased, more distinctions arose between the soil and wood microbes. At the genus level, the group Burkholderia-Caballeronia-Paraburkholderia presented as a unique, high-abundance genus in wood, averaging 13% of taxa in heartwood samples and 20.9% in sapwood samples. This genus is well recognized to be plant-associated ^60^, and has been documented for its ability to promote plant growth ^61^. Other unique, high abundance genera included *Conexibacter, 1174-901-12, Cellumonas, Methanobacterium, Oenococcus, Paenibacillus, Jatrophihabitans, and Sporolactobacillus* (LEfSe, p<0.01). All of these taxa have previously been reported to associate with plant tissues in some regard, being found on bark (*1174-901-1* ^62^*, Conexibacter* ^23^*, Sporolactobacillus* ^63^), roots (*Paenibacillus* ^64^), living and dead wood (*Cellumonas* ^65^*, Methanobacterium* ^66^*, Jatrophihabitans* ^67^), and even on oak barrels used for wine aging (*Oenococcus* ^68^). This evident enrichment of many known plant-associated taxa suggests that the microbiome of the wood of living trees is composed, at least in part, by an adapted community which could potentially have an active role in supporting or subverting tree health. For example, several taxa with biogeochemically-relevant metabolic capacities, including nitrogen fixers and methanogens– taxa themselves the subject of considerable interest as endophytes ^69–71^– were observed residing in the wood habitat (**Figure 2**).

For fungi, at the genus level, distinct taxa included *Pholiota*, a wood-rotting saprotroph ^72,73^, *Cladosporium*, a genus which hosts both plant pathogens and endophytes ^74^, and an unresolved genus in the family Teratosphaeriaceae, a group which includes stem and leaf pathogens ^75^. This collection of generally saprotrophic or pathotrophic fungal endophytes further corroborates class-level observations that the fungal microbiome within the wood of living trees may be well-situated to become pioneer decomposers upon tree death.

### Conservation of the Wood Microbiome at the Tree Species Level

We observed that the composition of microbial communities in the tissues of living trees varied significantly with tree species, suggesting that microbial communities within wood adapt and diversify in response to biochemical and physical variations among hosts (**Figure 2, 3**). Conditions within wood differ considerably amongst tree species, encompassing variations in pH, sap movement, phenolic levels, non-structural carbohydrate concentration, and the occurrence of antimicrobial agents, among others. PERMANOVA tests (Adonis2, WUnifrac, p<0.001) demonstrated that tree species was significantly related to both prokaryotic (heartwood: R^2^=0.31, sapwood: R^2^=0.33) and fungal community structures (heartwood: R^2^=0.44, sapwood: R^2^=0.49). Moreover, an association was observed between tree species and alpha diversity of both bacterial and fungal communities (p<0.05, ANOVA; **SI Figure 2**), suggesting variance in wood microbiome richness across tree species. Bacterial and fungal ordinations demonstrated this forest-wide variance, with many loose clusters forming along species lines (**Figure 2, 3**).

Moreover, the relatedness of wood-borne communities among tree species broadly followed the phylogenetic relationship among the trees themselves, suggesting specific microbiome adaptations to the host, and potential coevolution. Beta-diversity groupings (weighted Unifrac) of wood fungal communities correctly matched the phylogenetic clustering of all three birches (*Betula* spp.), both asterids (*Kalmia latifolia* and *Fraxinus americana*), and both gymnosperms (*Pinus strobus* and *Tsuga canadensis*) (**SI Figure 3**). For prokaryotic woodborne communities, phylogenetically-accurate clusters formed among all three birches, the maples, and oaks (**SI Figure 3**). As the likelihood of randomly observing three correct groupings in a perfectly bifurcating tree of 16 species is 1.18%, these groupings—seen repeatedly—are unlikely due to chance alone. This result supports microbial community adaptation to tree physiology or biochemistry, under the notion that more closely related tree species may have more similar microbial-habitat host conditions within their wood. This type of relatedness appears comparable to that found between closely related animals, such as seen in a study of non-human primates, wherein phylogenetic relatedness proved a key determiner of the gut microbiome, alongside environmental and dietary factors ^76^. As such, we may expect the wood microbiome to be shaped in large part by the environmental conditions within a host species. For instance, the maples (*Acer* spp.), which are recognized to contain high sugar content in their sap ^77^, presented with the highest heartwood concentrations (1.4 and 0.6%) of Saccharimonadia, a fermentative sugar-degrading group (**SI Figure 4**). Accordingly, we might expect the maple microbiome to broadly look similar across individuals given this biochemical uniqueness, but should two populations face differing conditions (e.g., onset of spring temperatures), we might expect some slight variances, at least temporally, in their microbiomes.

While the evolutionary history of trees may foster similarities in their wood microbiomes by creating more similar biochemical and physical conditions amongst closely related species, another possible mechanism for this appearance of relatedness could be a type of microbial inheritance ^78^. Such inheritance is a known process for animals and plants ^79^, and could involve the seeds of a tree carrying within them key taxa sourced from their parental microbiome. As trees speciated, divergence of inherited microbiomes along phylogenetic lines might be expected. This possibility of microbial inheritance raises further lines of inquiry about species level similarity in different stages of forest succession, and may even beg questions about the potential role of pollinators—a known plant microbiome vector ^80^—in shaping tree and wood microbiomes.

### The Wood Microbiome of Living Trees Appears Adapted to its Niche

Heartwood and sapwood prokaryotic communities were distinct from one another, with differentiation potentially driven by varying conditions within the two tissues, such as labile carbon and oxygen availabilities. Higher relative abundances of known anaerobic groups, like Clostridia, Negativicutes, and Methanobacteria, were observed in heartwood as compared to sapwood (Linear discriminant analysis Effect Size/LEfSe analysis, p<0.01) (**Figure 2**). Metabolic inference analyses (FAPROTAX) further demonstrated that anaerobic metabolisms, including fermentation, were also more abundant in the heartwood (**Figure 2**). Conversely, sapwood was enriched in obligate aerobes, such as Actinobacteria ^81^, with FAPROTAX inferring that aerobic chemoheterotrophy as a whole was more prevalent in sapwood (41.8%) than heartwood (28.2%). Conditions within heartwood are known to be broadly anaerobic ^82–84^ and nutrient-poor ^85^, whereas conditions within sapwood are more generally aerobic ^86^. This differentiation in conditions between tissues and their associated microbiota has previously been observed in some individual species ^87^, and may be further important for framing thought about the origin of woodborne taxa, as any potential inoculation of these regions would be expected to arise from environments with matching conditions (e.g., anaerobic). Alpha diversity was similar between the two sites (p>0.05), suggesting that community differences between tissues were not driven by varying levels of species richness.

Fungal community differences were also evident between heartwood and sapwood, but the distinctions were less pronounced than for prokaryotes. Tests (adonis2) confirmed that the amount of variance in community structure explained by tissue type was less for fungal (R^2^=0.019, p=0.003) than for prokaryotic communities (R^2^=0.047, p<0.001). This difference was visibly evident by a prominent overlap between fungal heartwood and sapwood communities in beta-diversity analyses (**Figure 3**). LEfSe tests further corroborated this finding, as, at the class level, there was only one fungal group unique to sapwood—lichen, or Lecanoromycetes. Lack of more dramatic fungal differentiation between heartwood and sapwood sites has been observed for the wood of individual species of living trees ^4^ and deadwood ^88–90^, suggesting that fungi may tolerate the differing conditions between tissues. This tolerance could be due to a potential ability of woodborne fungal taxa to switch between aerobic and anaerobic metabolisms, such as is the case for Saccharomycetes ^91^, a group observed herein. Alternatively, as fungal groups can build long hyphal networks, gas transport may afford them partial access to oxygen during colonization of anaerobic sites ^92^. Using tools to infer the functional and ecological roles of fungi (both FungalTraits, and see SI for FUNGuild), it emerged that saprotrophic and pathotrophic groups combined for nearly half of all resolved wood taxa, with symbiotrophic groups representing just 11-13% (**Figure 3**). These results imply a potentially more antagonistic role for woodborne fungal constituents, which could also explain why they appear less adapted to specific wood tissues, as their colonization may be more a function of opportunity than adaptation.

Surprisingly, despite the physical proximity of heartwood and sapwood, inferred source-sink movement between the sites accounted for a minority of the taxa appearing in either tissue, deepening questions about wood microbiome origins. For prokaryotic communities, source-tracking analyses (FEAST) estimated that the average transfer between heartwood and sapwood tissues was just 19%, with approximately equal rates of transfer in either direction (**SI Figure 5**). For fungal groups, this level of transfer was higher, at 29%, and there was greater contribution from the sapwood to heartwood (34%) than from the heartwood to sapwood (23%) (**SI Figure 5**). For prokaryotic communities, the heartwood had a higher proportion of taxa from unknown origin—alluding to either an uncharacterized source or endemic taxa—than sapwood (71% vs. 64%), whereas for fungal communities the opposite was observed (54% vs. 66%). This result is in alignment with previous observations suggesting that differences in conditions between sapwood and heartwood drive taxonomic differentiation through niche specialization. However, it also suggests that communities within heartwood and sapwood may have two distinct origin pathways, given their low level of overlap.

### Minimal Association of Soil Communities with the Wood Microbiome

Source-tracking analyses revealed a limited contribution of the soil microbiome to the wood microbiome, with estimates that only 3-5% of the community originated from mineral soils and 6-13% from organic soils (**SI Figure 5**). While expected ^93,94^ given the pronounced substrate and habitat differences, this result supports a relatively minor role of soil communities in shaping the wood microbiome. In our forest-scale survey, paired soil samples (from both organic and mineral horizons) were collected adjacent to each tree to explore soil origins of the wood microbiome, and whether variations in the wood microbiome were a function of varying soil communities. However, for all tissues, across both bacterial and fungal communities, source-tracking analyses (paired samples, FEAST) continually corroborated a modest contribution of the soil microbiome to the wood microbiome. The highest influence was observed in sapwood bacterial communities, wherein the combined contribution from organic and mineral soils peaked at 18%. As a large reservoir of microbial biodiversity, the soil might potentially still serve as a small dispersive source of taxa, but successful colonization of wood may be limited by the unique microenvironments within living trees. For instance, while soil taxa might try to migrate into trees, different predominating environmental conditions, including low oxygen and high CO_2_ concentrations within wood ^95^, may prohibit the infiltration of aerobic taxa from the soil ^96^, despite the known capacity of at least some pathogenic taxa to enter woody tissues from soil ^97^.

Across both prokaryotic and fungal taxa, the majority (54-71%) of taxa were ultimately found to have unknown sources—and thus are potentially endemic to their habitats. While soil is only one potential source, the fact that a large fraction of the wood microbiome appears unique to the interior of living trees suggests that the communities observed herein are unlikely to be just passive colonizers or a consequence of limited dispersion. Future work is required to unravel how dispersal, colonization and habitat conditions collectively determine the wood microbiome of living trees.

### The Wood Microbiome is Distinct Amongst Other Tree-Related Microbiomes

Intensive sampling of other neighboring microbial niches on, in, and around a focal tree revealed that the wood microbiome was distinct from communities found elsewhere. To further investigate community origins, we sequenced microbial DNA from a diversity of tissues (heartwood, sapwood, bark, branch wood, leaves, coarse and fine roots, and heart rot) and additional forest ecosystem compartments (mineral and organic soil, leaf litter) from a healthy, mature black oak (*Quercus velutina*) and its surrounding substrate. Despite this comprehensive survey (**Figures 4, 5**), heartwood samples still contained 104 unique (not found elsewhere) prokaryotic ASVs and 80 unique fungal ASVs, accounting for 43.2% and 21.6% of all heartwood sequences, respectively. Sapwood samples contained 91 unique bacterial ASVs and 1 unique fungal ASV, accounting for 8.7% and 0.1% of sapwood sequences, respectively. This high number of unique taxa appears more a result of niche specialization by wood-borne taxa, rather than a consequence of exceptional taxonomic diversity, as the alpha diversity measures for wood-borne communities, via both 16S and ITS, were on the lower-end of all sample types analyzed (**SI Figure 6**). This enduring differentiation of the wood microbiome, especially for heartwood, further corroborates the prior source-tracking conclusions that a large portion of woodborne taxa appears endemic to the interior of trees.

**Figure 4:**
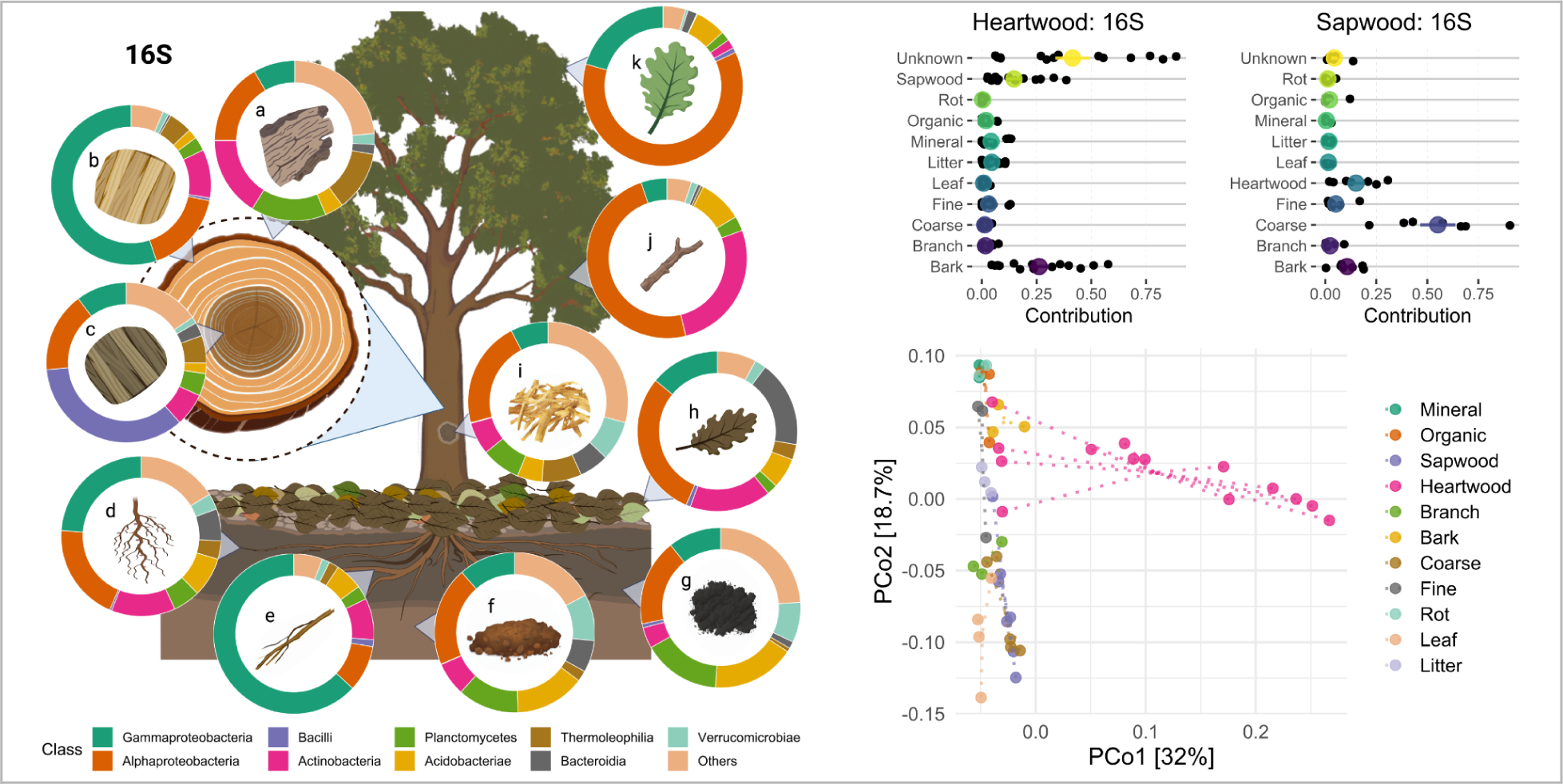
Overview of the black oak (*Quercus velutina*) prokaryotic microbiome. **a.)** Relative abundance of the top 9 prokaryotic classes (all other classes grouped in beige) in the a) bark, b) sapwood, c) heartwood, d) fine roots, e) coarse roots, f) mineral soil, g) organic soil, h) leaf litter, i) heart-rot, j) branches, and k) leaves. Source-tracking percent estimations (out of 1 or 100%) for microbial contribution from neighboring sites to the **b.)** heartwood and **c.)** sapwood microbiomes, based on FEAST analyses (taxa agglomerated at the species level). Mean value represented by the colored dot; SE represented by the bar. **d.)** Principal coordinate analysis for black oak tissues and surrounding environments, based on weighted UniFrac distance, with dashed lines converging on the centroid for each sample type.

**Figure 5:**
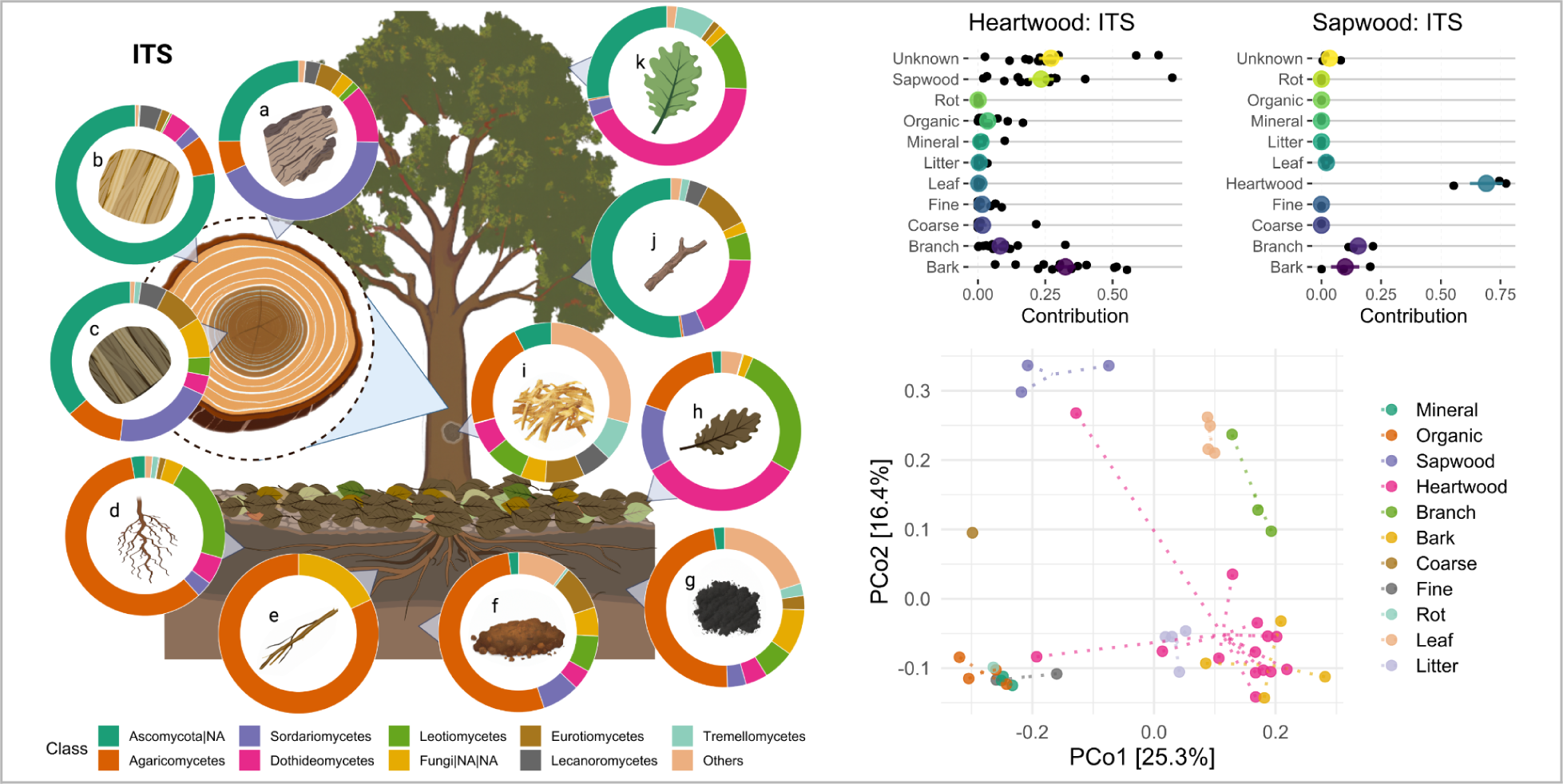
Overview of the black oak (*Quercus velutina*) fungal microbiome. **a.)** Relative abundance of the top 9 fungal classes (all other classes grouped in beige, with NA referring to unresolved groups; next resolved taxonomic level displayed) in the a) bark, b) sapwood, c) heartwood, d) fine roots, e) coarse roots, f) mineral soil, g) organic soil, h) leaf litter, i) heart-rot, j) branches, and k) leaves. Source-tracking percent estimations (out of 1 or 100%) for microbial contribution from neighboring sites to the **b.)** heartwood and **c.)** sapwood microbiomes, based on FEAST analyses (taxa agglomerated at the species level). Mean value represented by the colored dot; SE represented by the bar. **d.)** Principal coordinate analysis for black oak tissues and surrounding environments, based on weighted UniFrac distance, with dashed lines converging on the centroid for each sample type.

The microbial communities found in heartwood and sapwood proved more similar to other tree tissues than they did to one another, corroborating the marked differentiation between the two regions. Source-tracking analyses (FEAST using 16S data) suggested that almost half of the observed prokaryotic community in heartwood derived from unknown sources, with a smaller portion (25%) being associated with bark (**Figure 4**). For sapwood, however, the coarse root community was suggested to be the largest source, at over 50% contribution, with smaller contributions by heartwood and bark. For the fungal microbiome, similar results were found for heartwood (albeit with a diminished ‘unknown’ fraction), whereas sapwood was suggested to receive a majority of its contribution from heartwood (**Figure 5**). Supporting these results, 16S beta-diversity clusterings (**Figure 4**) found sapwood most similar to the tree’s coarse roots, whereas heartwood grouped by itself, between two groups of belowground (e.g., roots) and aboveground (e.g., leaves) samples. Conversely, for ITS clusterings, it was sapwood that grouped by itself, whereas heartwood grouped closely to bark (**Figure 5**).

These individual tree results broadly substantiate what was observed in the forest-level survey, such as the distinction between heartwood and sapwood communities, but they also introduce novel ideas about relatedness (**SI Figure 7**). For instance, the similarities observed between the communities of sapwood and other living tissues (i.e., coarse roots) may suggest that the communities within those sites are part of a larger living-wood microbiome, which is distributed throughout a tree by its interconnected hydraulic system. If true, this could potentially make variance in sapwood physiology a driver of community composition in living tissues. As trees can alter sap-flow in response to seasonal variation, age, and disease ^98^, community connectivity and similarity may shift in response to these conditions as well. Since resistance to hydraulic flow varies across the different architectures of a tree, like its roots and branches ^99^, differences in relative resistance to flow may help explain why regions still appear somewhat specialized despite overarching connectivity.

### Variance, or Lack Thereof, of Microbiomes with Height Reveal Further Tissue Differences

Multi-height sampling—from 50 cm to 10 m from the ground—revealed a sapwood microbiome that was stable with height, but a heartwood microbiome that varied significantly, casting the latter as a richer site of microbial diversity than previously realized (**SI Figure 8**). For the heartwood, which was sampled at seven different heights, virtually all (92%) of the variance in the prokaryotic microbiome across samples was associated with stem height, and more than half (56%) of the variance in the fungal community (p<0.01, adonis2, WUnifrac). This relationship with height was further apparent in beta-diversity analyses, especially so for the prokaryotic community, as samples neatly clustered accordingly (**SI Figure 8**). The composition of the sapwood community was not a function of height (p>0.05) using either 16S or ITS data. Conditions within sapwood, a living tissue, may generally be more regulated and homogenous, whereas for heartwood, conditions might vary more extensively with height due to gradients in tissue age, heterogeneities in wood structure (e.g., hollows, heart rot), or potential dispersal sources. While the uniqueness of the heartwood microbiome was suggested by the whole forest study, these individual-tree results suggest that future investigations may reveal unique communities across the vertical extent of a tree due to localized specialization of heartwood taxa. Conversely, the uniformity observed in sapwood microbial communities at different heights, along with their observed resemblance to taxa in other living tissues, further corroborates that these microbes may be part of a more extensive, stabler living-wood microbiome.

## Conclusions

Through this survey of bacterial, archaeal, and fungal taxa within the wood of over 150 trees across sixteen species, we have identified and described a distinctive microbiome resident within the wood of living trees, the largest pool of biomass on Earth. The wood of living trees is colonized by quantitatively substantial, seemingly adapted and specialized communities. These communities, potentially endemic to ecological niches within wood, may play significant roles in influencing tree health—a continuation of the recognized interactions between plants and their microbial associates ^100^. This research not only highlights a largely unexplored reservoir of biodiversity but also suggests potential opportunities for the discovery of novel and valuable compounds ^101^ and taxa with applications in ecosystem management, crop production, or human health and well-being. Uncovering the physiological and biochemical drivers of wood-microbiome diversity, and elucidating the consequences of microbiome composition and function for tree and ecosystem processes, may offer new avenues toward understanding the growth, management, and conservation of the world’s trees.

## Materials and Methods

### Study Site and Trees

All samples were collected from Yale-Myers Forest, a working and research forest in Northeastern, CT, USA (41.9529° N, 72.1239° W). The forest is classified in the Central Hardwood-Hemlock-Pine region^52^, and is composed mainly of second-growth vegetation which arose after colonial agricultural abandonment in the mid-19th century.

We selected fifteen dominant canopy trees in the region, as well as a dominant understory shrub, representing a wide range of species and wood traits, including both deciduous and evergreen species, hardwoods and softwoods, and different mycorrhizal associations. Species included: *Acer rubrum* (red maple, n=15), *Acer saccharum*, (sugar maple, n=5), *Betula alleghaniensis* (yellow birch, n=14), *Betula lenta* (black birch, n=20), *Betula papyrifera* (paper birch, n=6), *Carya ovata* (shagbark hickory, n=1), *Fagus grandifolia* (American beech, n=14), *Fraxinus americana* (white ash, n=12), *Kalmia latifolia* (mountain laurel, n=5), *Pinus strobus* (Eastern white pine, n=19), *Prunus serotina* (black cherry, n=3), *Quercus alba* (white oak, n=15), *Quercus rubra* (red oak, n=15), *Quercus velutina* (black oak, n=1), *Sassafras albidum* (sassafras, n=1), and *Tsuga canadensis* (Eastern hemlock, n=17). Excluding the shrub *Kalmia latifolia*, all trees sampled had a minimum diameter at breast height (DBH 1.37 m) of 51 cm, with no upper-bound on DBH. The median DBH across all species was 87.6 cm. These trees were sampled (following methods described below) during the peak growing season in 2021 (between July 19– August 12).

For the whole tree analysis, a mature black oak in a nearby mixed deciduous stand, also at the Yale-Myers Forest, was selected for sampling. While the tree had a visible seam in the bottom 1.5 m, indicating some potential heart rot, which was later confirmed, it appeared otherwise generally healthy, with a full canopy and minimal external signs of wounding or decay. The black oak was sampled following procedures described below at the end of the growing season in 2022 (October 4; prior to visible oak leaf senescence).

### Sample Collection

#### Wood Sampling

The wood microbiome was sampled through the collection of intact cores from standing live trees. Cores were extracted using 5.15-mm diameter increment borers (Haglöf Sweden, Sweden), which were decontaminated prior to each contact with a tree via flame-sterilizing with ethanol. Cores were sampled from tree surface to pith. Immediately following collection, cores were sealed in sterile aluminum foil and placed directly onto dry-ice until transfer back to Yale University, whereupon they were stored at −80°C until processing. Samples were collected during the peak growing season.

#### Soil Sampling

Soil samples were collected with a 5-cm diameter stainless steel soil probe, with a soil core taken from each of the four cardinal directions (N,S,E,W) around a given tree. Each core was collected at a distance to the tree equal to either the tree circumference, or 1 m, whichever was greater; this ensured the sample was taken beneath the tree’s canopy, within its rhizosphere, but in soil undisturbed by the root crown. Occasionally, if two trees were situated within a distance that meant soil samples would be taken within overlapping areas, or the environment prohibited sampling (e.g. the presence of a boulder or stream), the number of samples taken around each tree was reduced to three soil sampling points for each tree. For each core, the probe was inserted into the ground to 30-cm depth or the point of refusal. Upon retrieval, the soil cores were then separated into organic and mineral horizons. For each tree, the four mineral cores were bulked together to create a single composite mineral sample, and the four organic cores were bulked together to create a single composite organic sample. All composite soil samples were immediately passed through a 4-mm sieve in the field, and transferred into sterile 15-mL tubes. Soil samples were then immediately frozen on dry ice, and transferred to Yale University for storage at −80°C.

#### Black oak (*Quercus velutina*) sampling

For comprehensive sampling of the black oak, cores were collected from 50, 125, 200, 400, 600, 800, and 1000 cm from the ground. To enable concurrent sample collection, the tree was first felled using a chainsaw, with samples extracted immediately after. Additionally, samples of leaves, bark, branches, leaf litter, and heart rot were concurrently collected from the downed tree. To minimize contamination, tree-borne samples were only collected from areas not in contact with the ground. Additionally, samples of coarse (>2 cm) and fine roots (<2 cm) were collected from around the base of the tree by tracing and excavating roots connected to the stem. Soil samples were collected following the same procedure outlined above. All samples were collected in sterile plastic bags or centrifuge tubes, transferred to dry-ice immediately upon collection, and then transferred to −80°C for longer term storage.

#### Sample Processing and DNA Extractions

All wood cores were processed by first splitting them into corresponding segments of “heartwood” and “sapwood.” Heartwood was defined as the inner 5 cm of the core, as measured from the pith, whereas sapwood was defined as the outer 5 cm of the core, as measured from underneath the bark. Core segments were then lyophilized, from a frozen state and within individual sterile tubes with 0.22-μm filter caps, for 72 h, prior to cryo-grinding using a Spex 6775 Freezer/Mill® Cryogenic Grinder (Spex, NJ, USA). The grinding cycle was: pre-cool for 10 min, followed by 2 cycles of alternating grinding and cooling steps, each 2 min long. Grinding was performed at a rate of 10 cycles per second (CPS). Grinding vials were thoroughly decontaminated with bleach between each sample. DNA from the homogenized powder was then extracted using a MagMAX Microbiome Ultra Kit in concert with a KingFisher Apex automated extraction system. More details of this protocol, and critical validation, are presented in Arnold and Gewirtzman et al., 2024 ^35^. For the fine and coarse root samples taken from the black oak, roots were first washed and debrided in DI water prior to processing. Leaves, roots, bark, litter, and heart rot samples were then lyophilized and cryo-ground the same as for the wood core samples.

For DNA extractions from soil, 250 mg of soil (wet) were used for extraction via the MagMAX Microbiome Kit in concert with the Kingfisher Apex. Prior to proceeding with the extraction, samples and supplied lysis buffer were bead-beat for 10 min at maximum speed using a Vortex Genie 2 outfitted with a Vortex Adapter. The manufacturer’s protocol and program (MagMAX_Microbiome_Soil_Liquid_Buccal_v1.kfx) were followed, with final elution into 100 μL of buffer.

### Sequencing and Quantification

#### UMGC Sequencing

Extracted DNA was sent to the University of Minnesota Genomics Center (UMGC) for ITS (internal transcribed spacer) and 16S rRNA amplicon sequencing. For ITS sequencing, primers (5.8SR_Nextera, ITS4_Nextera) targeting the ITS2 region were used, and for 16S sequencing, primers (515F/806R) targeting the V4 region of the 16S rRNA gene were selected. These universal primers target fungal and prokaryotic (bacterial and archaeal) groups, respectively. In order to limit amplification of plastid DNA from tree tissue, both mitochondrial and chloroplast peptide nucleic acid (PNA) clamps were used during library construction (PNA Bio, Newbury Park, CA, USA). For sequencing, libraries were dual-indexed and sequenced on an Illumina MiSeq platform using a paired-end flow cell employing v3 (2 x 300bp) chemistry.

#### qPCR

Quantitative polymerase chain reaction measurements of 16S abundances (using universal V4 primers) were performed by the UMGC using the extracted DNA sent for sequencing. For each reaction, 3 μL of extracted DNA was used, with the number of cycles set to 35. A standards curve spanning 7 orders of magnitude (10-10,000,000 copies per reaction) was run concurrently with the samples, with R^2^ values >0.9.

### Bioinformatic Analyses

#### Pre-processing

Nephele v2.23.2 ^102^, using the DADA2 ^103^ R package v1.28, was used for pre-processing of sequence data, including filtering, trimming, merging of paired-end sequences, chimera removal, and taxonomic assignment. For 16S sequencing, forward reads were truncated to 260 bp, reverse reads were truncated to 180 bp, maxEE was set to 5, truncQ was set to 4, and the SILVA ^104^ v138.1 database was used for taxonomic assignment. The median read count for 16S data following pre-processing was 10,014 reads, with 86.7% of ASVs resolved at the class level. For ITS data, primers were removed using cutadapt v4.5, maxEE was set to 5, truncQ was set to 4, min_len was set to 50, and taxonomic assignment was done using the UNITE v8.3 database. The median read count for ITS data following pre-processing was 21,957 reads, with 52.8% of ASVs resolved at the class level.

#### Post-processing

Following pre-processing, sequence data was imported into R v.4.3.0 for analysis using phyloseq ^105^ v.1.44.0 and microeco ^106^ v.0.17.0. An unrooted phylogenetic tree was used for ITS data, while a rooted tree was used for 16S data. For ITS sequence data, samples were rarefied to a depth of 4,000 reads (**SI Figure 14**). For 16S sequence data, remnant chloroplast and mitochondrial ASVs were first filtered out (10.3% of ASVs), and then samples were rarefied to a slightly lower depth (3,500 reads) to account for sequence loss during plastid filtering. Rarefaction curves were produced by Nephele using plot.ly v.2.5.1. For source-tracking analyses, FEAST ^107^ v.0.10 was used. All analyses, unless otherwise specified, were paired (at the level of the individual tree), with Expectation-Maximization iterations set to 10,000. Taxa were agglomerated at the family level for the whole-forest study, whereas they were agglomerated at the genus level for the black oak study. For construction of the phylogeny of tree species, the phytotools ^108^ package v.1.5-1 and S.PhyloMaker function were employed, drawing on the PhytoPhylo megaphylogeny as a backbone. For maps of tree genera distribution, the rnaturalearth ^109^ v.0.3.3 package was used for illustration of data sourced from the GlobalTreeSearch database ^110^. For broad-level ordinations (e.g., between different sample types), dimensional reductions were performed using principal component analyses (PCoAs) on weighted unifrac distance. For within-sample-type ordinations, where differences between sample communities were expected to be finer, dimensional reduction was performed on unweighted unifrac distances using uniform manifold approximation and projection (UMAP) via the umap v.0.2.10 package ^111^. For differential abundance analyses, the LEfSe method was employed using microeco. For LEfSe analyses, the significance level was set to 0.01, and the threshold was set to 3.5. Taxa unresolved at the class level were removed. In the case of functional inference, the FAPROTAX v.1.2.6 database ^112^ was utilized for prokaryotic communities, whereas the FungalTraits ^113^ and FUNGuild ^114^ databases were used for fungal communities. For heatmaps based on functional inference, weighted abundances were used. Adonis2 PerMANOVA tests ^115,116^ were performed within microeco.

### Data Availability

The sequencing data generated during this study will be available in the NCBI database upon publication. Code and additional datasets supporting the findings of this study will be made available in FigShare prior to publication.

## Acknowledgements

We extend our gratitude to Michelle Spicer, Joseph Orefice, Matt Valido, Thomas Harris, Mark Ashton, Ben Girgenti, Laura Logozzo, Taylor Maavara, Judith Rosentreter, Les Welker, Alex Polussa, Makenzie Birkey, Marsh Hlavka, Ellie Jose, Camila Ledezma, Ari Gewirtzman, and Cyrena Thibodeau for their invaluable assistance in the field. We also thank Claire Butler, Adriana Rubenstein, Morgan Furze, Darryl Angel, Jonas Karosas, Talia Kolodkin, and Eli Ward for assistance in the laboratory. We acknowledge the University of Minnesota Genomics Center for its technical assistance. Thank you to the Yale Forests faculty, staff, and facilities for enabling the research.

Funding for this research was provided by the National Science Foundation Graduate Research Fellowship Program (NSF-GRFP), the Yale Institute for Biospheric Studies (YIBS), and the Kohlberg-Donohoe Research Fellowship to Jonathan Gewirtzman; the National Defense Science and Engineering Graduate (NDSEG) Fellowship to Wyatt Arnold; and additional support from the Yale Center for Natural Carbon Capture and the Yale Planetary Solutions Project to Jordan Peccia, Mark A. Bradford, Peter A. Raymond, Craig Brodersen, Marlyse Duguid, Jonathan Gewirtzman, and Wyatt Arnold.

## Supporting Information

### Supplemental Methods

#### Estimated Cell Quantification

As is broadly reported for the human microbiome, the ratio of human cells to bacterial cells is roughly 1:1^36,117^. In a cubic centimeter of wood, the number of plant cells may vary from around 180,000 to 5,700,000, depending on cell size^118^. Assuming an average wood density of 0.5g cm^−3^ and a wood moisture content of 50%, we can estimate the number of plant cells in a dry gram of wood as between 720,000-22,800,000, or roughly 10,000,000 on average. Based on the prokaryotic abundances we observed in wood—averaging 10^6^.^35^ copies per gram—and assuming four 16S copies per average cell^45^, we assert that there is approximately 1 bacterial cell for every 20 plant cells within a gram of dry wood. Given that 16S abundances vary considerably by tree species, this ratio would shift in accordance, from as low as 1:600 in *Prunus serotina* to as high as 1:2 in *Acer saccharum*. Assuming a tree weighing approximately 5000 kg (wet), this would put the number of prokaryotes in the wood of a standard living tree at around 1 trillion cells, or 1/38^th^ the number inside the average human^36^. When taking into consideration communities in the rhizosphere, phyllosphere, and dermosphere, the number of microbes inhabiting the whole of the tree biomass will be much greater.

## Supplemental Figures and/or Extended Data Figures

**SI Figure 1:**
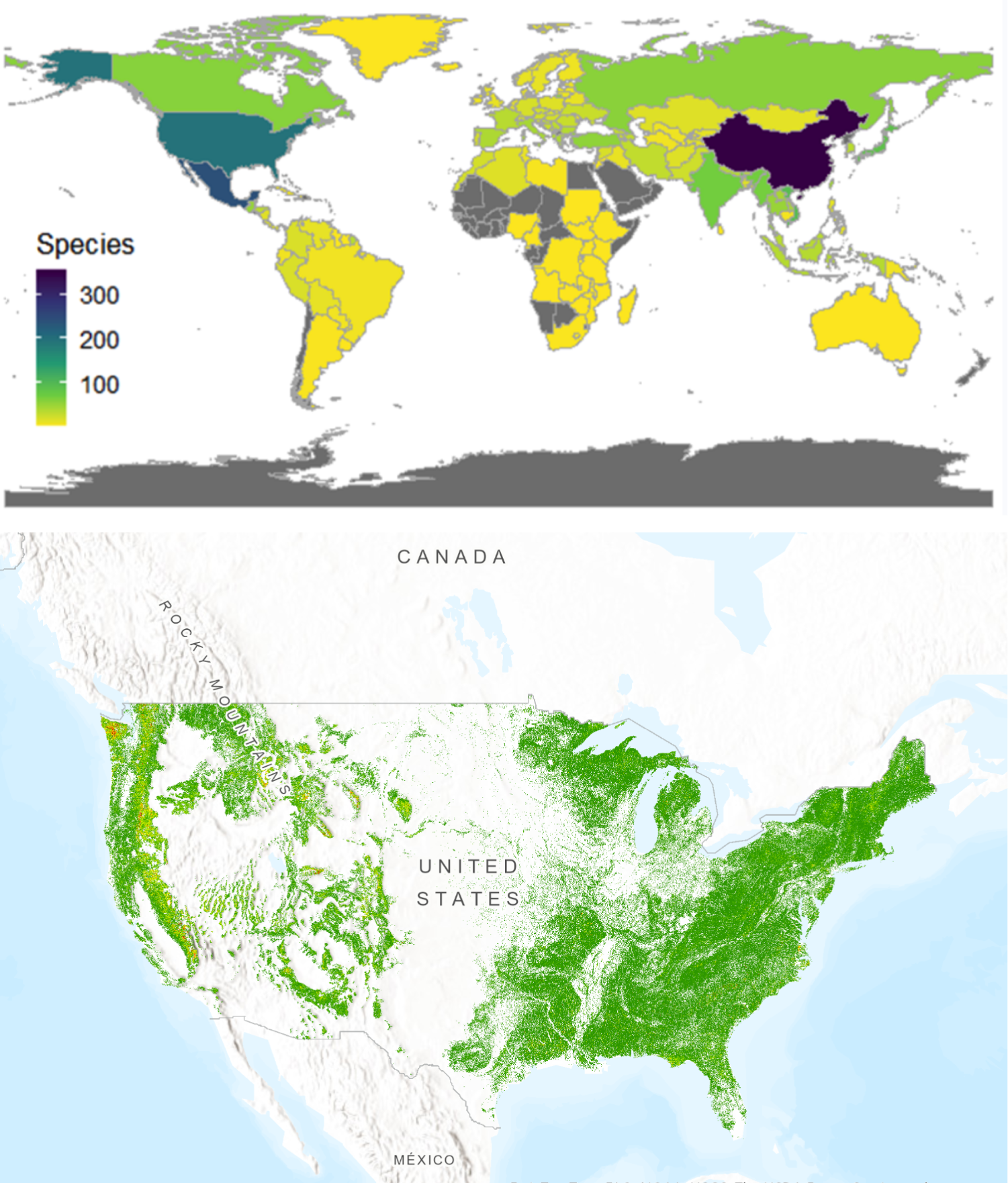
The species included in this survey represent 11 genera which have **A**) a global distribution, with the map colorized by the number of species within this set of genera native to each country. Within **B**) the United States, the genera studied herein have a distribution that spans much of the East Coast and Midwest, and extends sporadically through to the West Coast (from US Forest Service Individual Tree Species Parameter Maps).

**SI Figure 2:**
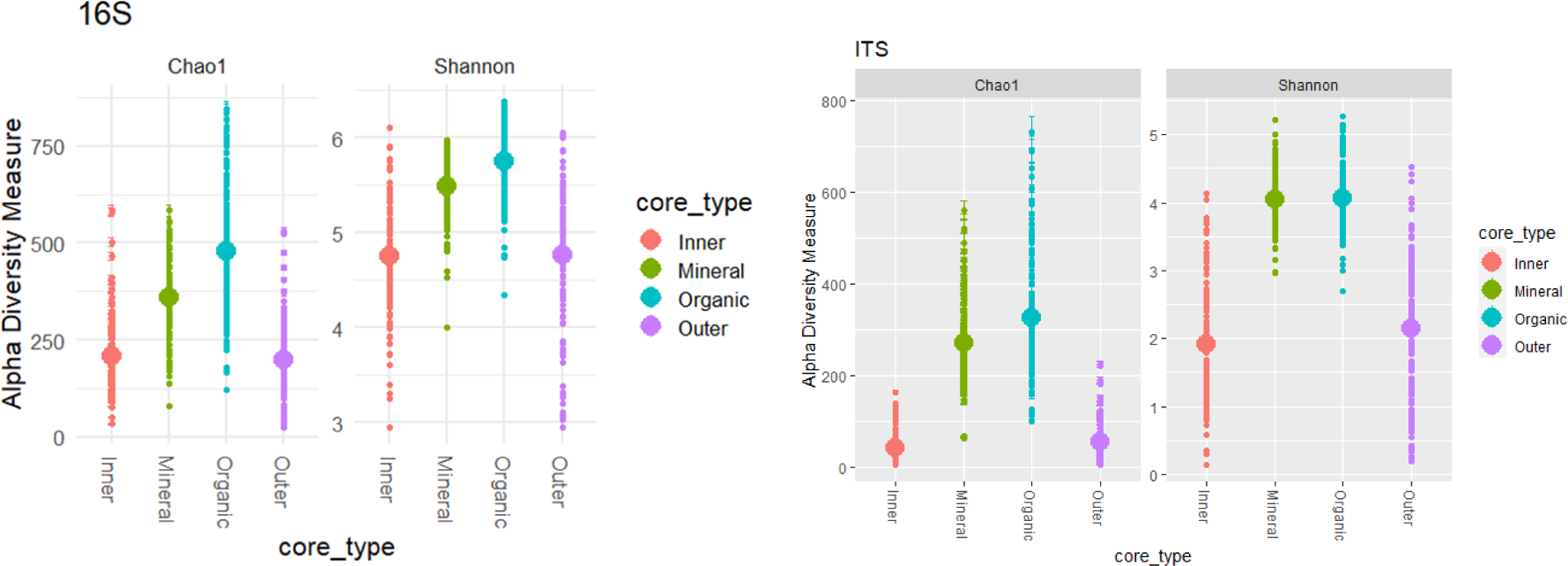
Alpha diversity (using Chao1 and Shannon metrics) for the whole-forest study, split by sample type (Inner=heartwood, Outer=sapwood, Mineral=mineral soil, Organic=organic soil) and amplicon (16S and ITS). Mean value for measure displayed via the larger point.

**SI Figure 3:**
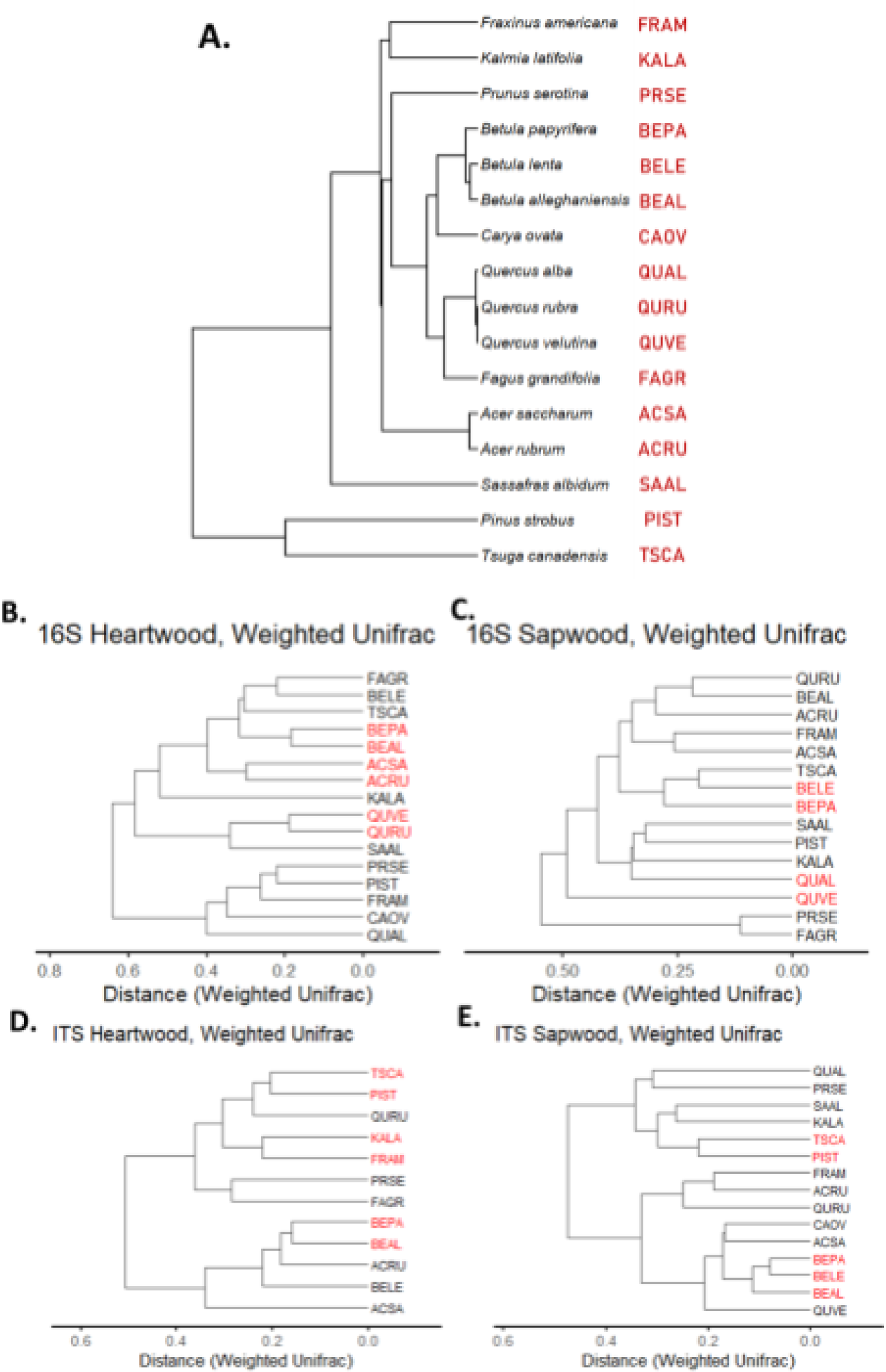
The **A.**) phylogenetic relatedness of the 16 tree species included in this study, based on the PhytoPhylo megaphylogeny. Both Latin binomial names and species codes are included at the tips. Using both **B, C.)** 16S sequencing data and **D, E.**) ITS sequencing data, phylogenies showing the relatedness of living wood microbiomes between tree species were produced based on beta diversity (weighted Unifrac distance) similarity. Species codes in red represent tip pairings between microbial communities that match the phylogenetic relatedness between host tree species.

**SI Figure 4:**
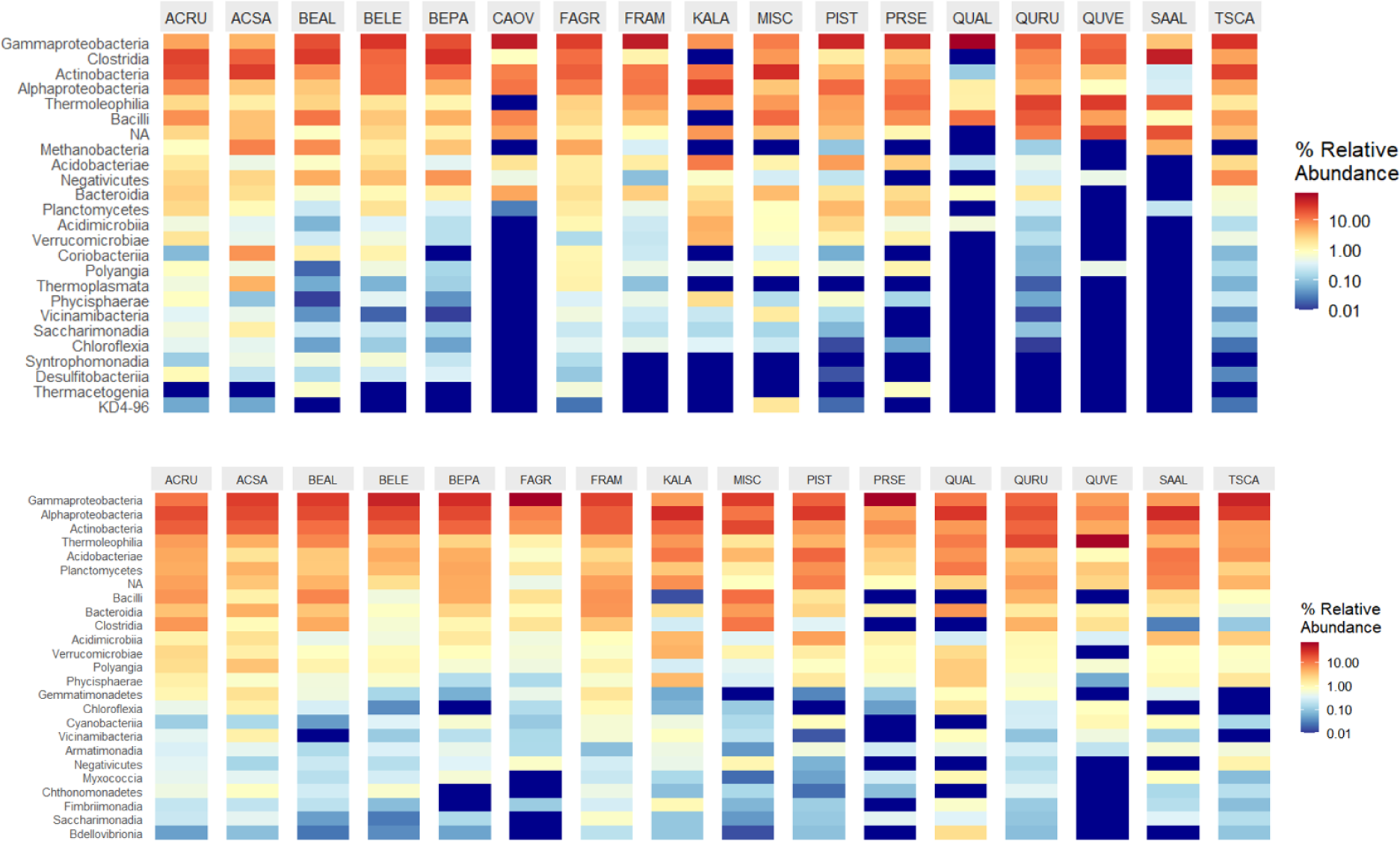
Relative abundances of 16S taxa at the class level for both **a.)** heartwood and **b.)** sapwood samples, split by tree species (see SI Table 1 for codes).

**SI Figure 5:**
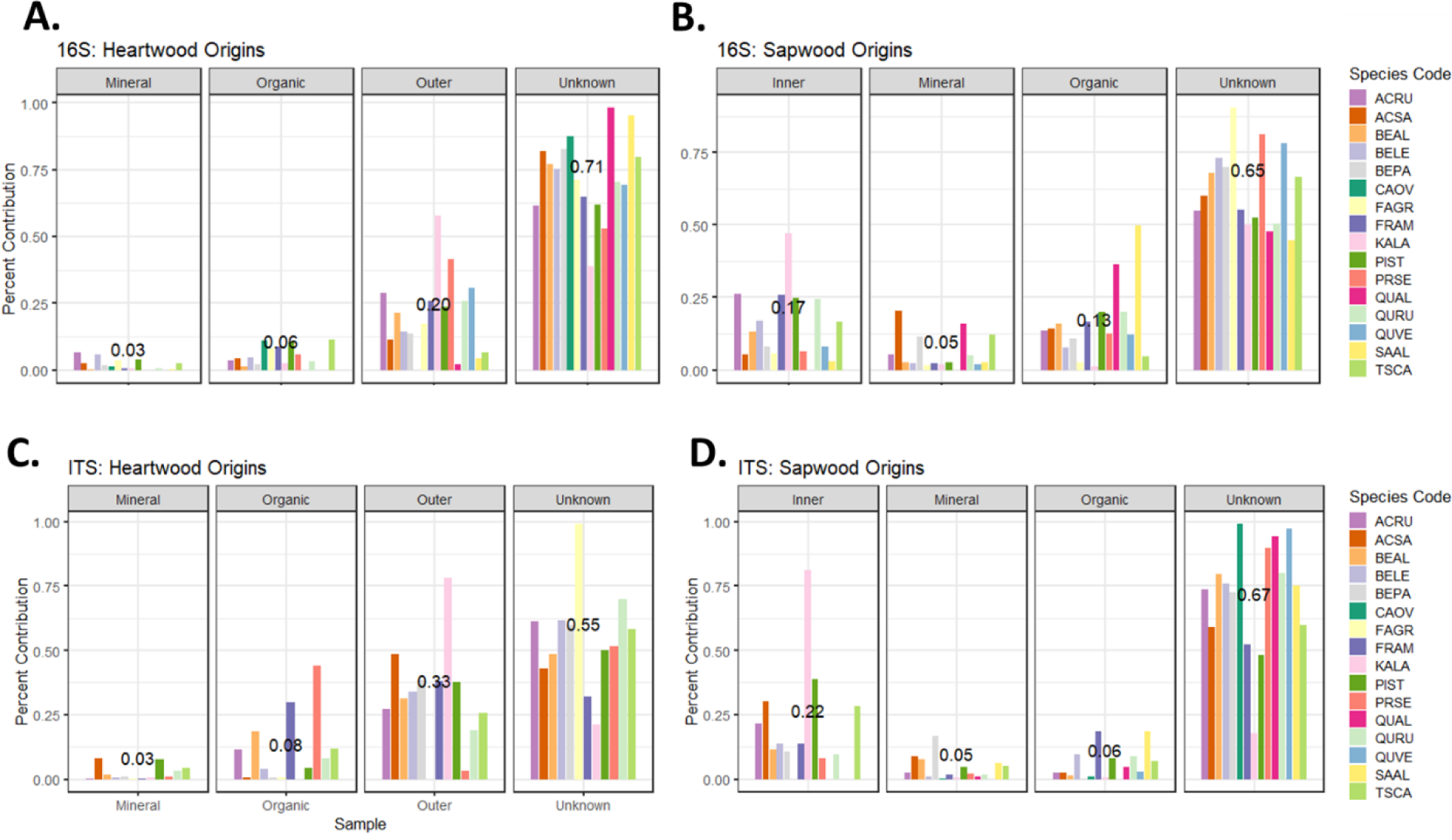
Source-tracking results produced using FEAST, treating both **A, C**.) heartwood and **B, D**.) sapwood as “sinks.” For heartwood analyses, the contribution of sapwood, organic soil, and mineral soil communities was assessed, with “unknown” corresponding to taxa with indeterminate origins. For sapwood, the contribution of heartwood, organic soil, and mineral soil communities was assessed. Results are split by tree species, with the central number representing the average contribution out of all samples (out of 1). Analyses were performed in a paired manner, meaning that contribution estimates were assessed on a per tree basis (e.g., wood and soil samples specific to each tree, rather than bulk groups).

**SI Figure 6:**
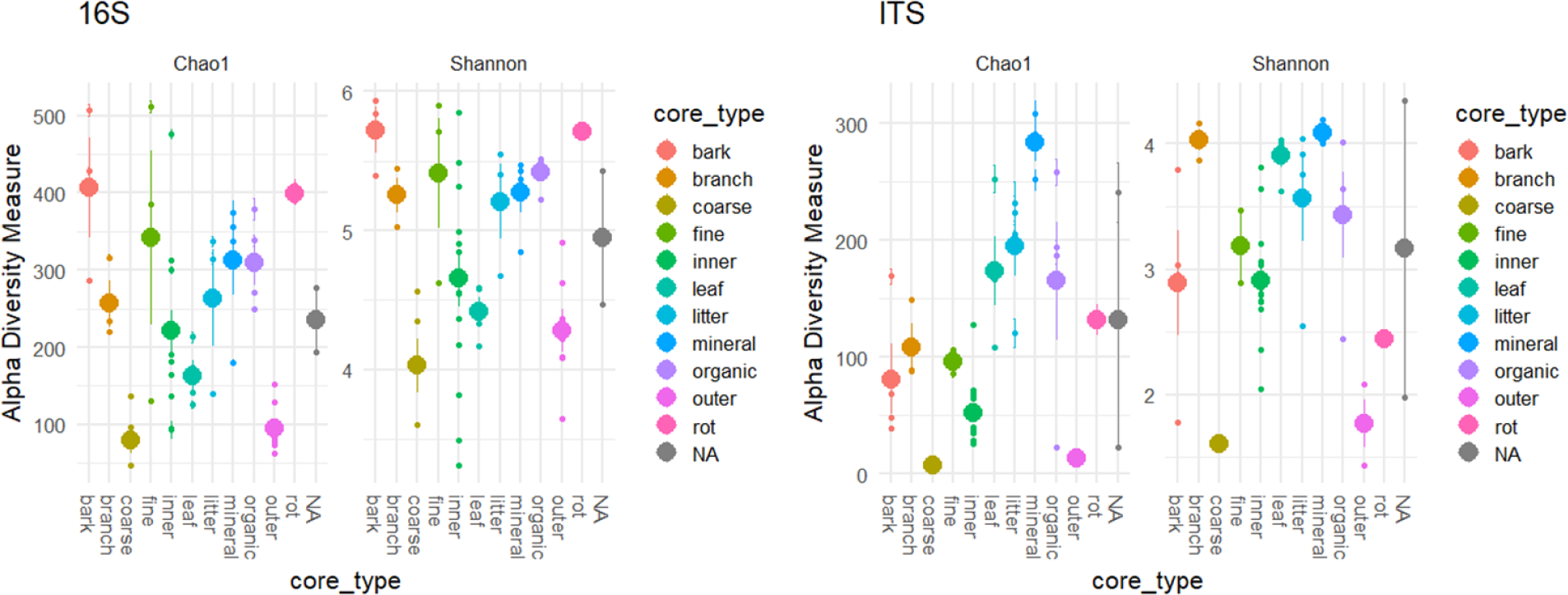
Alpha diversity estimates, using Chao1 and Shannon indices, for varying tissue and environmental samples from the black oak study, split by **A**.) prokaryotic and **B**.) fungal communities. Large dots represent mean diversity, with the line representing ± SE.

**SI Figure 7:**
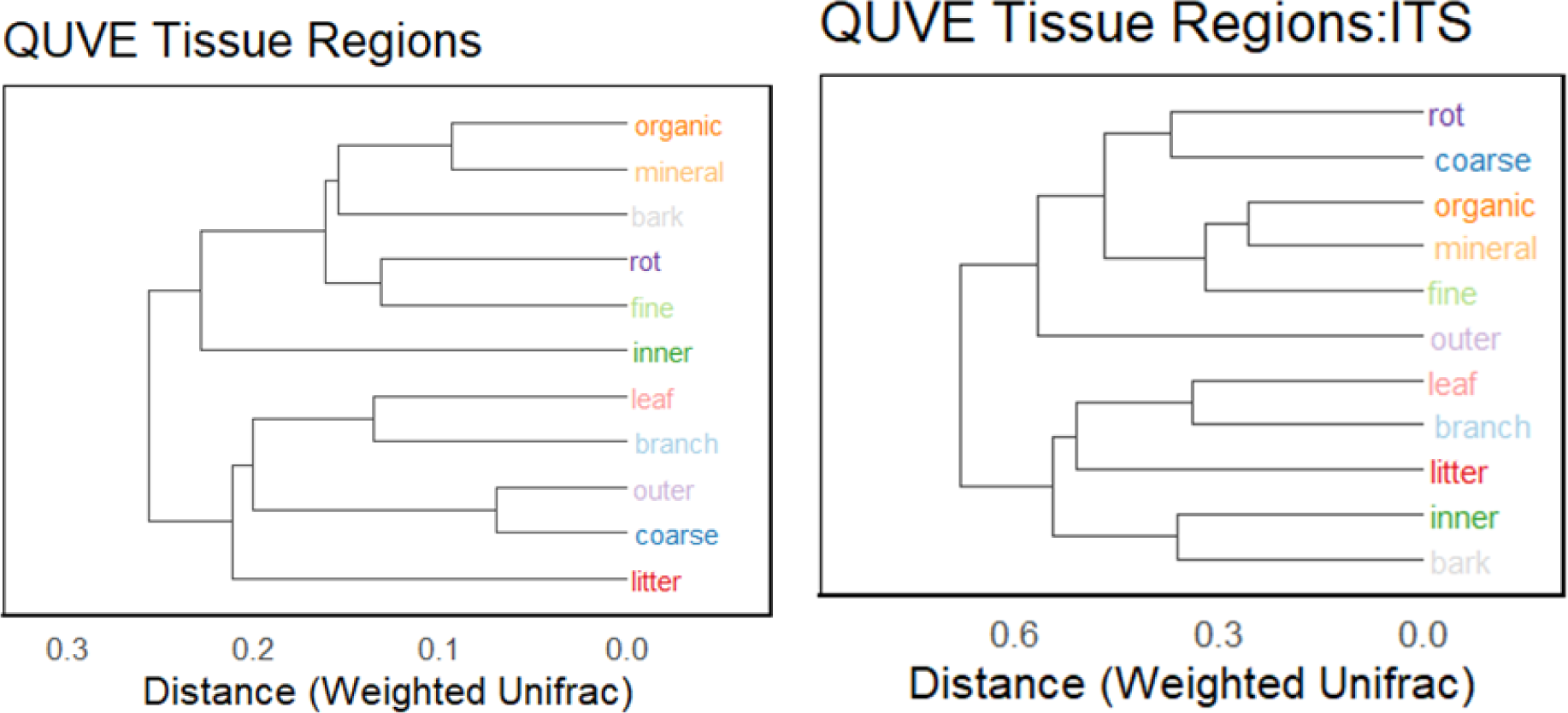
Relatedness of samples to one another **B, E**), based on weighted Unifrac distance (coarse and fine refer to roots, rot refers to heart-rot).

**SI Figure 8:**
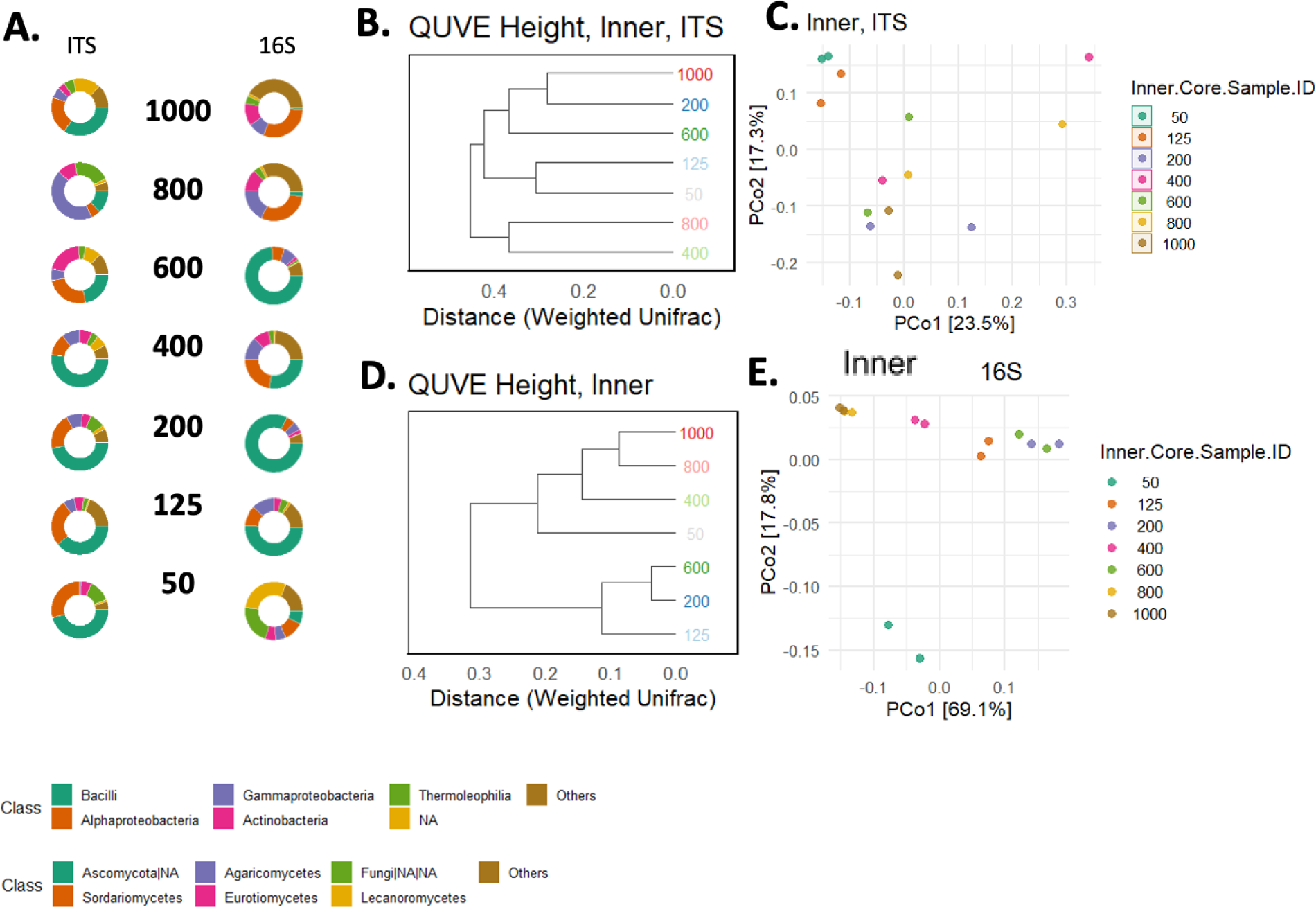
Variation in the **A**. fungal and prokaryotic heartwood microbiomes with tree height in the black oak (central numbers represent distance from ground in cm). The relatedness between those tissues, based on weighted Unifrac distances, is represented in **B.** and **D**., with ordination (PCoA, weighted Unifrac) of those same tissues in **C.** and **E**.

**SI Figure 9:**
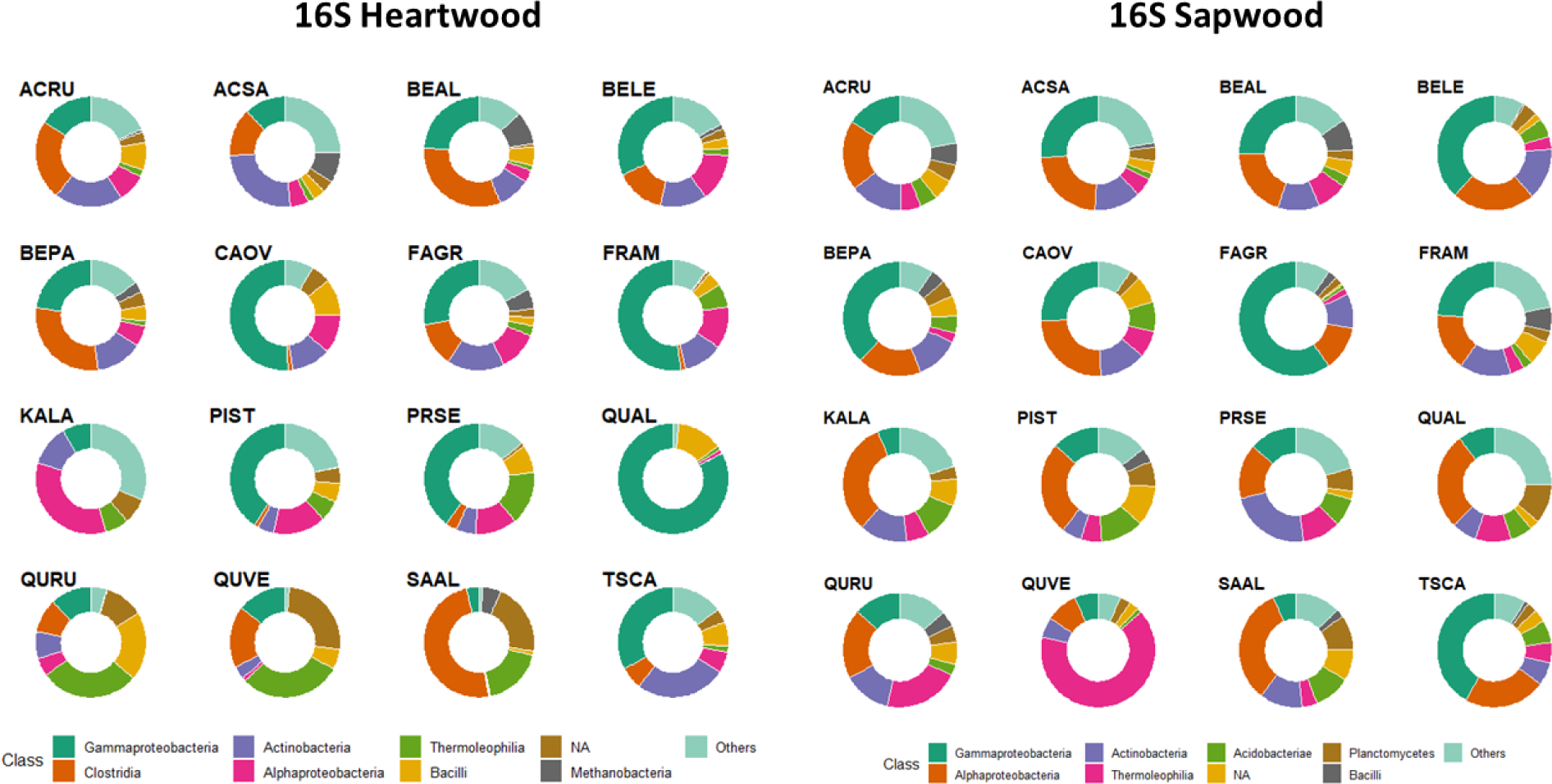
Average prokaryotic relative abundances at the class-level, split by heartwood and sapwood, for the 16 species included in this study. Only the first 8 most abundant classes are shown (remaining grouped into “Others”).

**SI Figure 10:**
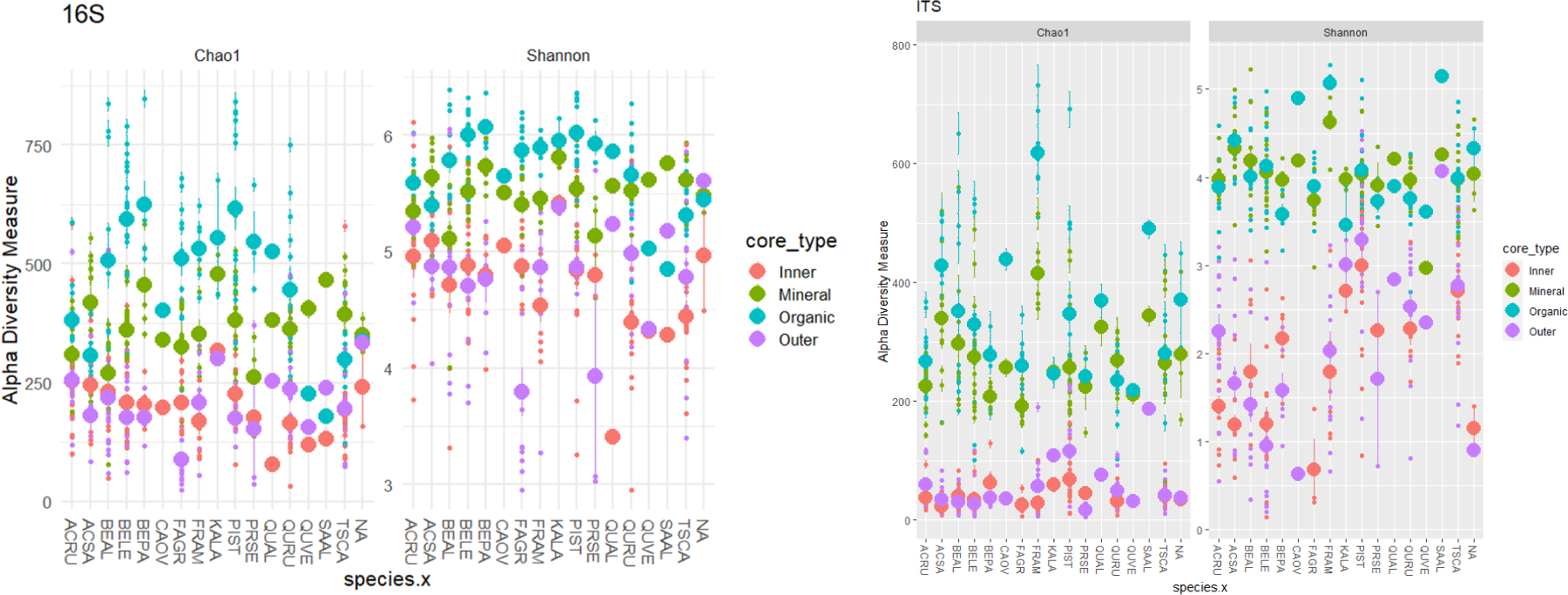
Alpha diversity measures (Chao1 and Shannon) for **a.)** 16S and **b.)** ITS communities, split by tree species (“species.x”) and sample type (“core_type”; Inner=heartwood, Outer=sapwood, Mineral=mineral soil, Organic=organic soil). Mean (± SE) represented by the larger point and underlying line.

**SI Figure 11:**
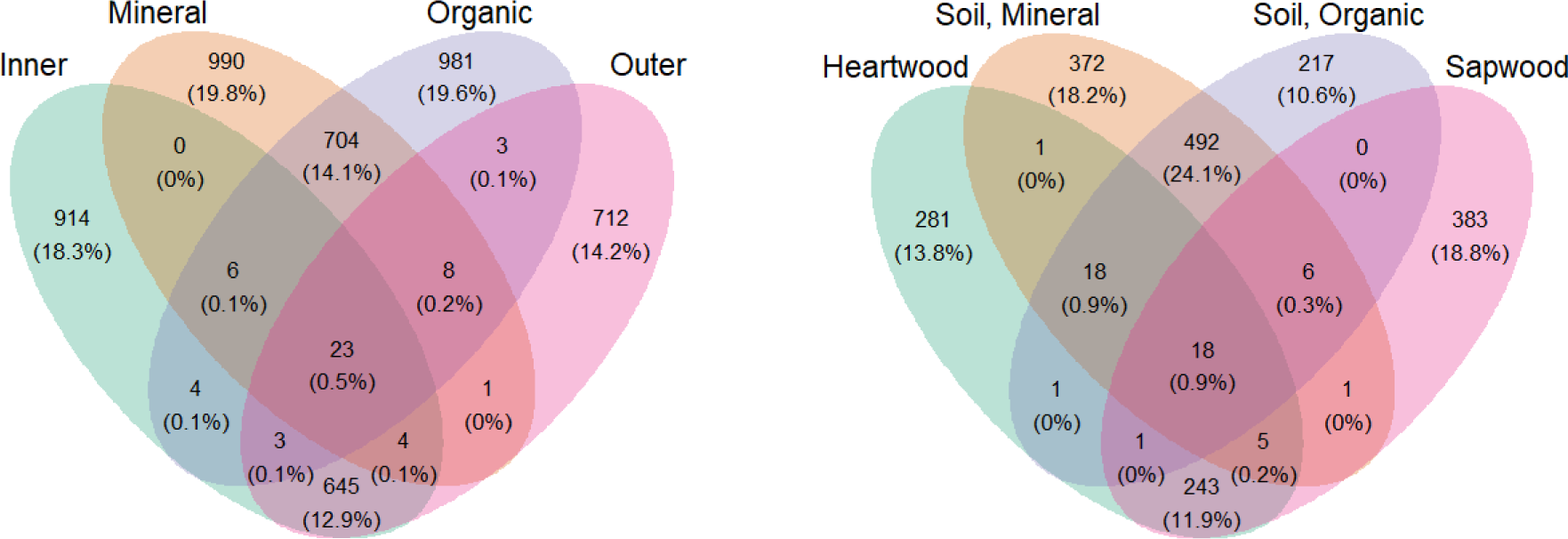
16S (left) and ITS (right) counts of unique or shared taxa in each forest compartment. Percent represents percentage of total ASVs.

**SI Figure 12:**
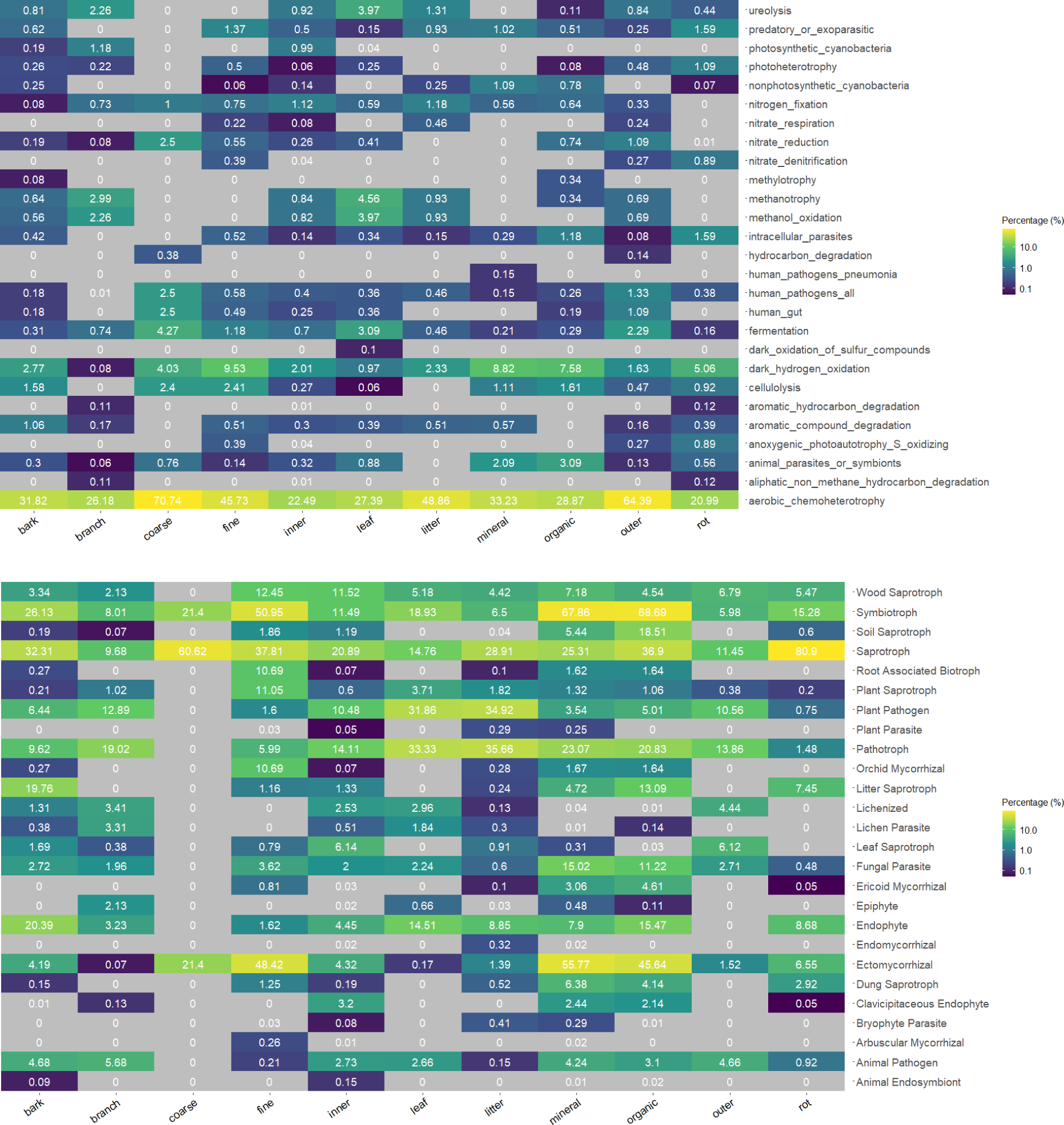
Inferred **a.)** 16S metabolisms using FAPROTAX and **b.)** fungal lifestyles using FungalTraits for black oak samples, split by sample type.

**SI Figure 13:**
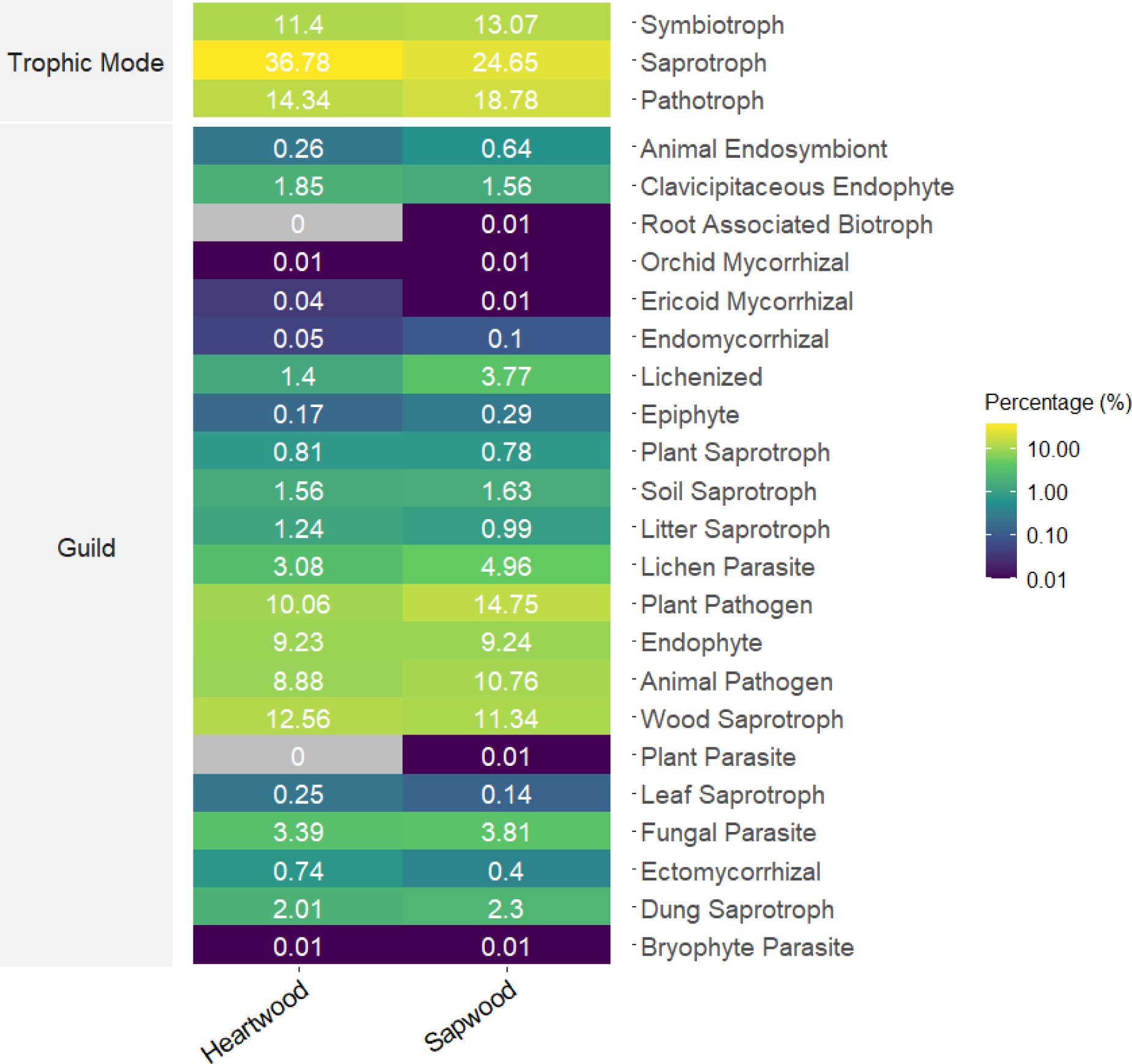
Percent abundance (out of 100) of fungal trophic modes and guilds within heartwood and sapwood samples, as inferred by FUNGuild.

**SI Figure 14:**
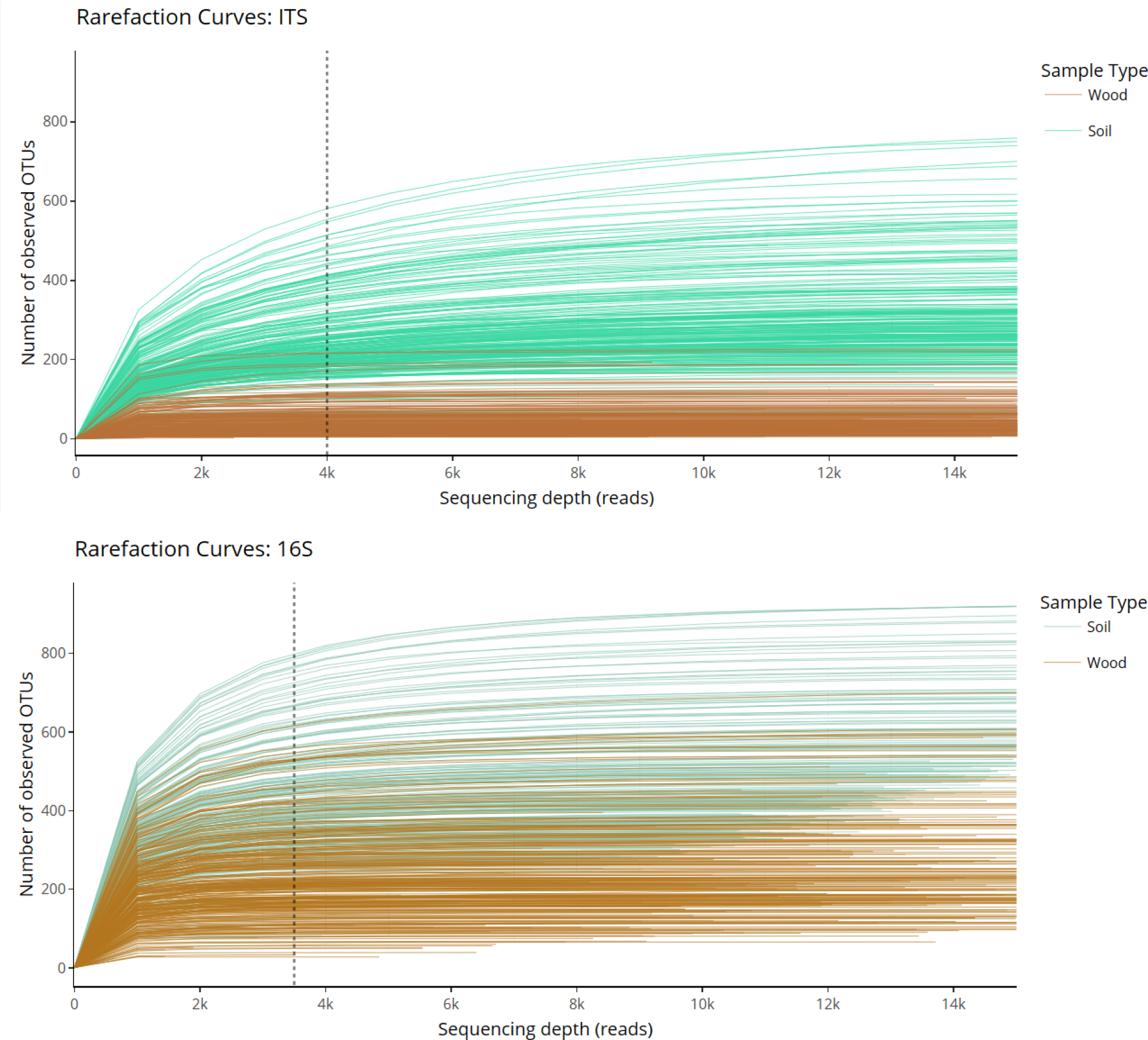
Rarefaction curves for **a.)** ITS and **b.)** 16S sequencing data. Wood samples displayed in brown, with soil samples displayed in teal. For ITS data, samples were rarefied to 4,000 reads per sample (dotted line), and for 16S data, samples were rarefied to 3,500 reads per sample (dotted line) after filtering out remnant mitochondrial and chloroplast sequences.

## Supplemental Tables

**SI Table 1:**
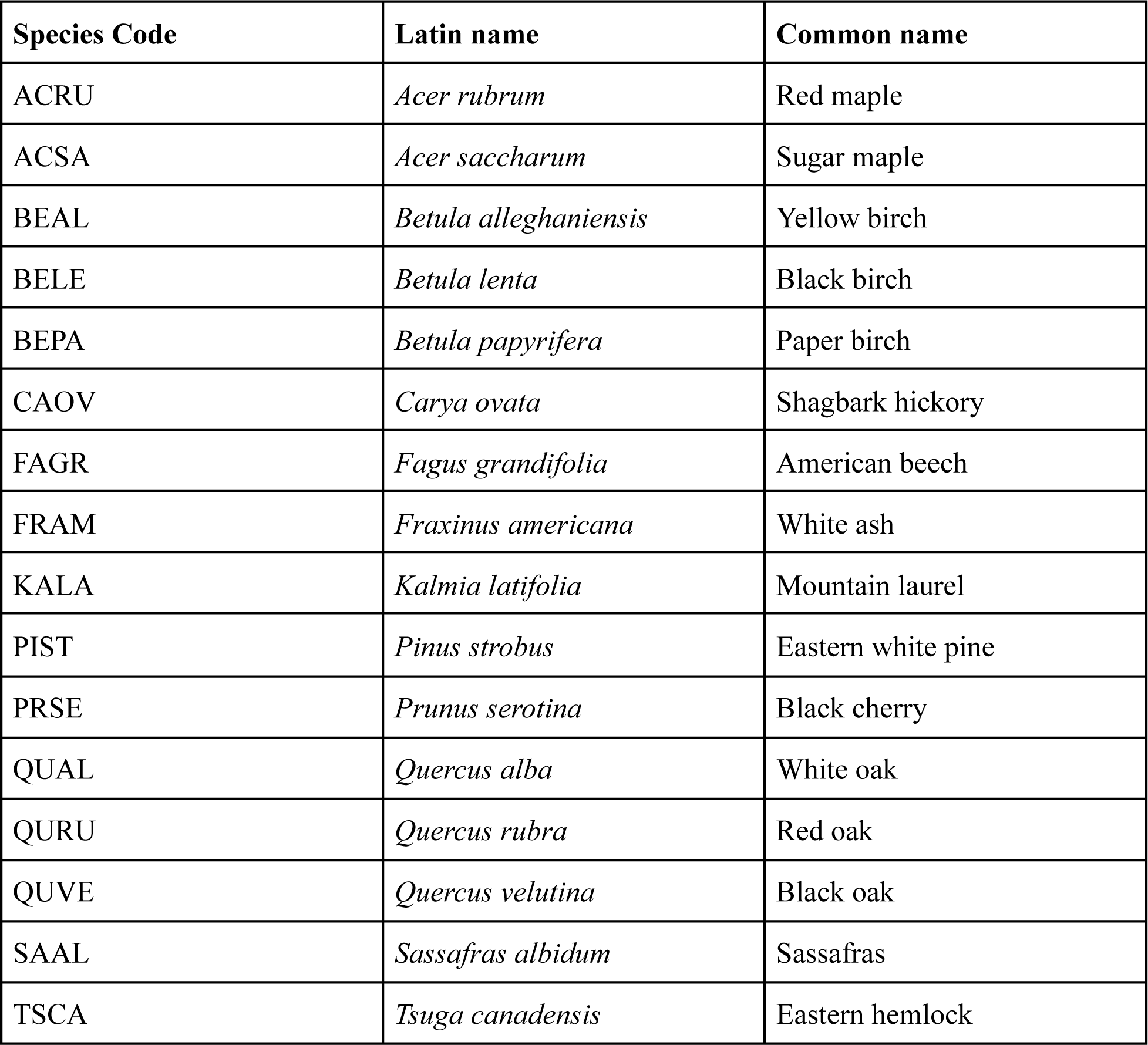
Species codes, scientific names, and common names for species in this study.

| Species Code | Latin name | Common name |
| --- | --- | --- |
| ACRU | *Acer rubrum* | Red maple |
| ACSA | *Acer saccharum* | Sugar maple |
| BEAL | *Betula alleghaniensis* | Yellow birch |
| BELE | *Betula lenta* | Black birch |
| BEPA | *Betula papyrifera* | Paper birch |
| CAOV | *Carya ovata* | Shagbark hickory |
| FAGR | *Fagus grandifolia* | American beech |
| FRAM | *Fraxinus americana* | White ash |
| KALA | *Kalmia latifolia* | Mountain laurel |
| PIST | *Pinus strobus* | Eastern white pine |
| PRSE | *Prunus serotina* | Black cherry |
| QUAL | *Quercus alba* | White oak |
| QURU | *Quercus rubra* | Red oak |
| QUVE | *Quercus velutina* | Black oak |
| SAAL | *Sassafras albidum* | Sassafras |
| TSCA | *Tsuga canadensis* | Eastern hemlock |

